# NRF2-driven thiol metabolism creates a selective vulnerability to selenite

**DOI:** 10.64898/2026.11.12.766318

**Authors:** Ting Chao, Yun Huang, Niklas B. Thompson, Jing-Chun Lin, Ryan Martin, Rithwik Paladugu, Yi-Han Tang, Svitlana Altukhova, Shayla Pham, Basma A. Yasseen, Feroza K. Choudhury, Archi Patel, Min Liu, Bin Fang, John Koomen, Jose Serrano-Velez, W. Gregory Sawyer, Hardik Shah, Mingxiang Teng, Chang Jiang

## Abstract

NRF2 hyperactivation caused by mutations in *KEAP1* or *NFE2L2* occurs in 20-30% of non-small cell lung cancers and confers therapy resistance. Here we show that NRF2-driven metabolic reprogramming creates a selective vulnerability to selenite. NRF2-dependent thiol export promotes the extracellular reduction of selenite to selenide. Our data support a model in which selenide enters cells and reaches mitochondria, where sulfide-quinone oxidoreductase uses selenide and glutathione to generate the selenopersulfide GS-SeH. A four-hour pulse of selenite was sufficient to induce cell death after washout, consistent with a selenium redox cycle that consumes NADPH and regenerates selenide. Within hours, selenite abolished aconitase activity and impaired mitochondrial respiration. The resulting cell death occurred independently of apoptosis, ferroptosis, pyroptosis, necroptosis and disulfidptosis. In mice, a selenite-rich diet selectively suppressed the growth of NRF2-hyperactive tumors. These findings identify an NRF2-driven metabolic vulnerability and support further investigation of selenite as a therapeutic strategy for NRF2-hyperactive cancers.

---

Mutations in the KEAP1-NRF2 pathway occur in approximately 18% of lung adenocarcinomas (LUAD) and 25-29% of lung squamous cell carcinomas (LUSC)^1,2^. Under physiological conditions, NRF2 is continuously targeted for ubiquitination and proteasomal degradation by the KEAP1-CUL3 ubiquitin ligase complex. Disruption of KEAP1 stabilizes NRF2, allowing its accumulation in the nucleus and activation of antioxidant response genes. Persistent NRF2 activation confers resistance to chemotherapy, radiotherapy and immunotherapy^3–8^. Targeting this pathway could therefore benefit a substantial proportion of patients with NSCLC, yet effective therapeutic strategies for KEAP1-NRF2-mutant tumors remain lacking.

NRF2 lacks a catalytic domain, making this transcription factor difficult to target directly. High-throughput screens have identified NRF2 inhibitors, but their utility has been limited by poor selectivity or toxicity^9^. More recently, VVD-065 was developed to promote NRF2 degradation by binding Cys151 of wild-type KEAP1 and enhancing KEAP1-CUL3 complex formation. This compound suppresses NRF2-dependent tumor growth, restores sensitivity to chemotherapy and radiotherapy, and has entered a phase I trial (NCT05954312)^10^. However, its requirement for an intact KEAP1-NRF2 interaction limits its applicability to tumors harboring mutations that disrupt this interaction. Thus, approaches that suppress NRF2 independently of KEAP1 remain necessary. An alternative is to exploit the metabolic state established by NRF2 activation. NRF2-hyperactive cells increase cystine uptake through xCT and maintain elevated glutathione synthesis^11,12^. Compounds whose toxicity depends on this thiol-rich state could therefore selectively target these treatment-resistant tumors.

Selenium is an essential micronutrient incorporated into 25 human selenoproteins as selenocysteine, the 21^st^ amino acid, in which selenium replaces the sulfur atom of cysteine. Eleven selenoproteins contribute to antioxidant defense, including the glutathione peroxidases and thioredoxin reductases^13–18^. The chemical properties of selenocysteine, including its low pKa, support efficient redox catalysis and resistance to irreversible oxidation^19,20^. These functions have motivated investigations of selenium in cancer prevention and treatment and in other diseases, although trials of dietary selenium supplementation have yielded largely inconclusive results^21–32^. Dietary selenium is available in organic forms, predominantly selenomethionine, and inorganic forms, including selenite and selenate, which are commonly used in supplements. Following absorption and metabolism, selenium is distributed to tissues in part by selenoprotein P (SELENOP), a circulating selenium transport protein secreted primarily by the liver^33,34,35^.

Inorganic selenium differs chemically from the selenomethionine used in those trials and can exert cytotoxic effects independently of selenoprotein synthesis. Selenite reacts readily with thiols through a series of glutathione-dependent reactions involving selenodiglutathione and the selenopersulfide GS-SeH, while producing selenide^36–38^. These reactions oxidize thiols and generate reactive intermediates, establishing selenite as a redox-active cytotoxin^39^. At concentrations above the nutritional range selenite is toxic to cancer cells and has been evaluated in two phase I trials. Intravenous sodium selenite was tolerated at doses up to 10.2 mg/m^2^ in 34 patients with chemotherapy-refractory carcinoma^40^, whereas oral sodium selenite was tolerated at doses up to 33 mg daily alongside palliative radiotherapy in patients with metastatic disease^41^. Cellular selenium uptake from selenite depends on extracellular reduction by cysteine, supported by cystine import through xCT, a mechanism proposed to underlie its selective toxicity towards chemoresistant cancer cells^42^. Selenide also serves as a substrate for mitochondrial sulfide-quinone oxidoreductase (SQOR), which transfers electrons from selenide to ubiquinone. The resulting ubiquinol limits lipid peroxidation and suppresses ferroptosis^43^.

Together, these findings implicate uptake rate as a determinant of selenite selectivity and reveal a protective role for intracellular selenium metabolism. However, what determines whether selenium is tolerated or becomes lethal remains unclear, and the genetic alterations that confer selenite sensitivity have not been defined. NRF2 provides a potential link between these questions because it regulates the transporter and thiol pools that govern selenite uptake and metabolism.

Here we show that NRF2 hyperactivation creates a selective vulnerability to selenite. NRF2-driven thiol export promotes extracellular reduction of selenite to selenide, which is metabolized in mitochondria by SQOR to form the glutathione selenopersulfide GS-SeH. We confirm the identity of this species using a synthetic standard. Our findings support a redox cycle in which GS-SeH formation and resolution consume NADPH while regenerating selenide. This cycle maintains the chemically available selenium level inside the cell. Selenite disrupts mitochondrial iron-sulfur cluster function and impairs respiration within hours, inducing cell death independently of apoptosis, ferroptosis, pyroptosis, necroptosis, and disulfidptosis. In mice, a selenite-rich diet suppresses the growth of NRF2-hyperactive tumors without affecting KEAP1-intact tumors.

## NRF2 hyperactivation promotes selenium accumulation and sensitizes NSCLC cells to selenite-induced cell death

To determine whether NRF2 status influences selenite sensitivity, we profiled a panel of NSCLC cell lines with wild-type (WT) or altered *KEAP1/NFE2L2*. Cell lines harboring KEAP1 mutations or NFE2L2 amplification expressed elevated levels of NRF2 protein and its downstream targets, xCT^44^, GCLC, GCLM^45,46^, distinguishing NRF2-high from NRF2-low groups (Fig. 1a). Across the panel, NRF2-high cell lines exhibited significantly lower sodium selenite (Na_2_SeO_3_) IC_50_ values than NRF2-low cell lines (Fig. 1b).

**Figure 1:**
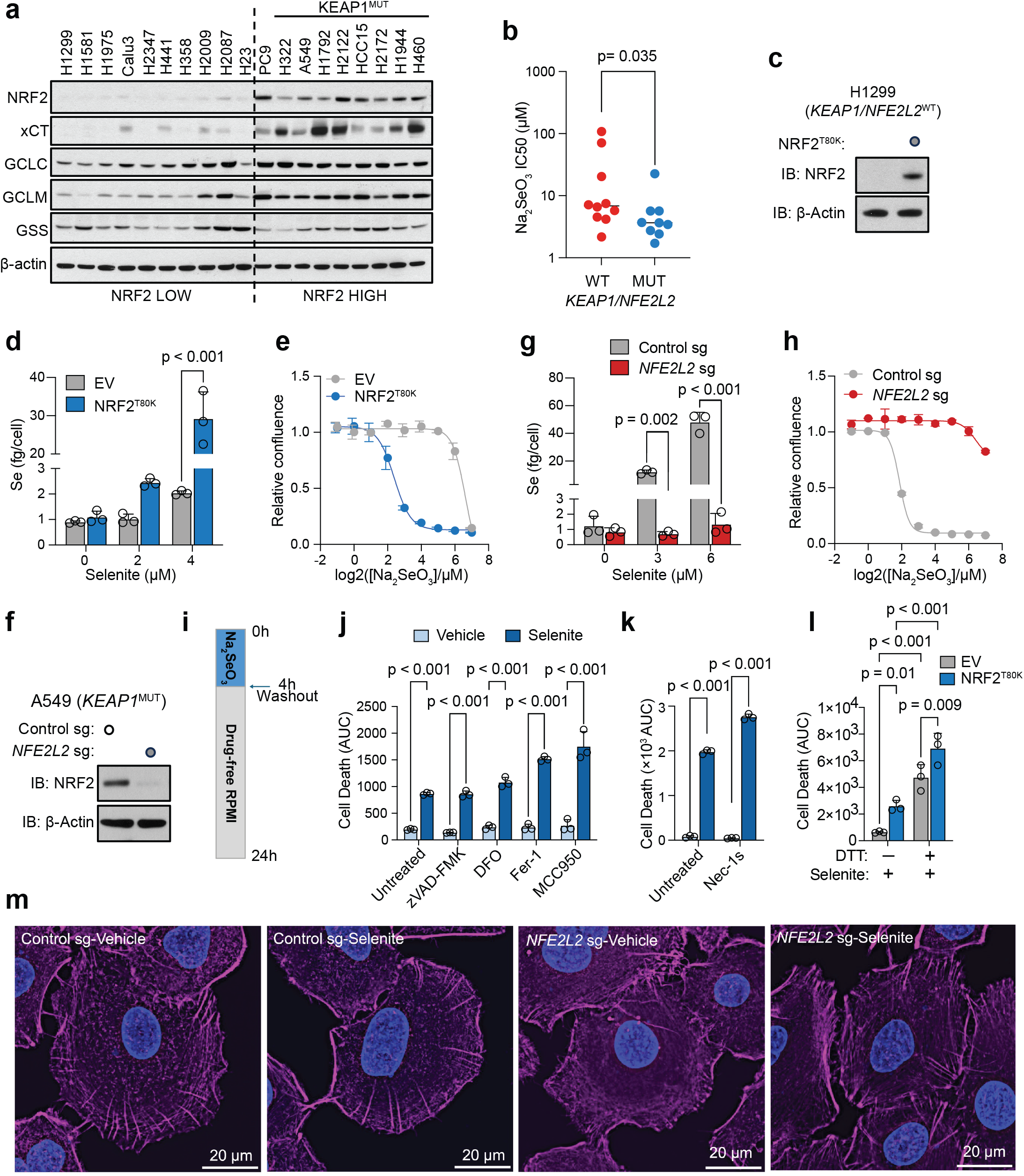
NRF2 hyperactivation sensitizes NSCLC cells to selenite through a non-canonical cell death mechanism. **a**. Immunoblots of NRF2, GSS and NRF2 target proteins xCT, GCLC, GCLM across a panel of NSCLC cell lines grouped by *KEAP1/NRF2* genotype; β-actin, loading control. **b**. Na_2_SeO_3_ IC50 values for the lines listed in **a**, grouped by *KEAP1/NFE2L2* status. IC50 values were log_10_ transformed prior to Mann-Whitney test analysis; n = 10 (WT), n = 9 (MUT) cell lines. **c**. Immunoblot of NRF2 in H1299 cells transduced with empty vector (EV) or NRF2^T80K^. **d**. Intracellular selenium of the cells depicted in **c**. after 4 h exposure to the indicated Na_2_SeO_3_ concentrations, measured by ICP-MS via ^78^Se. **e**. Dose-response of the cells depicted in **c**. to listed Na_2_SeO_3_ concentrations, measured as relative confluence by live-cell imaging. **f**. Immunoblot of NRF2 in A549 cells (*KEAP1*-mutant) transduced with a non-targeting control sg or *NFE2L2* sgRNA. **g**. Intracellular selenium in the cells depicted in **f** after 4 h exposure to 0, 3, and 6 µM Na_2_SeO_3_. **h**. Dose-response of the cells depicted in **f** to Na_2_SeO_3_. **i**. Schematic illustration for selenite treatment: cells were exposed to Na_2_SeO_3_ for 4 h, washed off, and returned to drug-free RPMI + 10% FBS for the remainder of the experiment. **j**. Cell death in H1299 EV and NRF2^T80K^ cells, quantified as the area under the curve (AUC) of SYTOX Green positive cell counts normalized to total cell area during 24 hours of the experiment as described in **i**. zVAD-FMK (25 µM), DFO (10 µM), ferrostatin-1 (2.5 µM), or MCC950 (15 µM) were added 18 h before the experiment, concurrently treated with or without 8 µM Na_2_SeO_3_ for 4 h, and re-supplemented for the next 20 h after Na_2_SeO_3_ washout. **k**. Cell death assayed as in **j** with or without necrostatin-1s (25 µM) **l**. Cell death assayed as in **j** with or without DTT (500 µM, added concurrently with 18 h before Na_2_SeO_3_). **m**. Representative confocal microscopy images of A549 cells transduced with control sgRNA or *NFE2L2* sgRNA, stained for F-actin with phalloidin (magenta) and nuclei with DAPI (blue), post exposure to vehicle or 8 µM Na_2_SeO_3_ for 4 h. Scale bars, 20 µm. Data in **b, d, e, g, h** and **j-l** are presented as mean ± s.d. Statistical significance was assessed by Mann-Whitney test; or two-way ANOVA followed by Tukey’s multiple-comparison test.

Next we used isogenic cell models to test whether NRF2 activity accounts for this difference. Expression of NRF2^T80K^, a mutant resistant to KEAP1-mediated degradation, in *KEAP1/NFE2L2*-WT H1299 cells (Fig. 1c) increased intracellular selenium accumulation in a dose-dependent manner following sodium selenite exposure (Fig. 1d) and lowered the concentration required to inhibit proliferation (Fig. 1e). Time-course measurements revealed a sharp increase in selenium accumulation between 2 and 4 h in NRF2^T80K^-expressing cells, whereas empty-vector (EV) controls showed a delayed and gradual increase over 12 h (Extended Data Fig. 1a). Conversely, CRISPR-mediated depletion of *NFE2L2* in *KEAP1*-mutant A549 cells (Fig. 1f) reduced selenium accumulation (Fig. 1g) and conferred resistance to sodium selenite across the concentrations tested (Fig. 1h).

Similarly, overexpression of WT *NFE2L2* in the *KEAP1/NFE2L2*-WT cell lines H1581 and H23 increased selenium accumulation and shifted the selenite dose-response curves towards lower concentrations (Extended Data Fig. 1b-g), indicating that elevated NRF2 is sufficient to confer sensitivity. Conversely, *NFE2L2* deletion in the *KEAP1*-mutant cell line HCC15 conferred resistance (Extended Data Fig. 1h, i). This phenotype was conserved in murine cells: *Keap1* disruption in KP (*Kras*^LSL-G12D/+;^ *Trp53*^fl/fl^) lung tumor cells increased NRF2 protein levels and sensitized cells to selenite (Extended Data Fig. 1j, k). Together these findings establish NRF2 activity as a determinant of cellular selenium accumulation and sensitivity to selenite.

We next investigated the cell death pathways involved in this response. Because cellular selenium accumulation plateaued by 4 h (Extended Data Fig. 1a), we used a pulse-washout regimen in which cells were exposed to sodium selenite for 4 h, then returned to selenite-free standard medium (Fig. 1i). Cell death continued after selenite removal. The pan-caspase inhibitor zVAD-FMK, the iron chelator deferoxamine (DFO), the ferroptosis inhibitor ferrostatin-1, the NLRP3 inflammasome inhibitor MCC950 (Fig. 1j), and the RIPK1 inhibitor necrostatin-1s (Fig. 1k) all failed to rescue selenite-induced cell death.

Similar results were obtained in A549 cells, in which liproxstatin-1 also failed to rescue cytotoxicity (Extended Data Fig. 2b). SYTOX Green staining revealed extensive loss of membrane integrity in selenite-treated A549 control cells, which was absent in *NFE2L2*-knockout (KO) cells (Extended Data Fig. 2a). These results suggest that selenite-induced cell death does not depend on caspase-mediated apoptosis, ferroptosis, NLRP3-dependent pyroptosis or RIPK1-dependent necroptosis.

Because NRF2-hyperactive cells express elevated levels of xCT (Fig. 1a), we also considered disulfidptosis. This is a form of cell death associated with cystine-dependent NADPH depletion, aberrant disulfide crosslinking of actin cytoskeletal proteins and F-actin collapse^47,48^. Two observations argue against this mechanism. First, whereas thiol-reducing agents suppress disulfidptosis, dithiothreitol (DTT) enhanced selenite-induced cell death in both EV and NRF2^T80K^-expressing cells (Fig. 1l). Second, phalloidin staining after 4 h of selenite exposure revealed an intact F-actin network, without the actin contraction or detachment from the plasma membrane characteristic of disulfidptosis (Fig. 1m). Moreover, all experiments were conducted in glucose-replete medium without glucose starvation or inhibition of glucose uptake, conditions that typically induce disulfidptosis. Together, these findings indicate that selenite-induced cell death in NRF2-hyperactive cells is distinct from disulfidptosis.

NRF2-dependent sensitivity extended to other inorganic selenium compounds. Sodium selenide (Na_2_Se), selenium dioxide (SeO_2_) and selenous acid (H_2_SeO_3_) each induced cell death at lower concentrations in NRF2^T80K^-expressing H1299 cells than in EV controls. Conversely, *NFE2L2* disruption in A549 cells conferred significant resistance to all three compounds (Extended Data Fig. 2c-e).

## NRF2 promotes extracellular thiol-dependent reduction of selenite

The NRF2 target xCT exchanges intracellular glutamate for extracellular cystine and supports the release of reduced thiols into the medium, suggesting that NRF2 may promote extracellular selenite reduction before cellular uptake. Expression of *SLC7A11*, which encodes xCT, in *KEAP1*-wild-type H1299 cells conferred selenite sensitivity (Fig. 2a, b). Conversely, *SLC7A11* disruption in A549 cells abolished xCT expression and conferred resistance to sodium selenite (Fig. 2c, d). Pharmacological inhibition of xCT with erastin protected cells across the selenite concentrations tested (Fig. 2e) and reduced selenium accumulation in NRF2^T80K^-expressing cells to near-background levels (Extended Data Fig. 3a, b). Similarly, *SLC7A11* disruption prevented selenium accumulation in A549 cells (Extended Data Fig. 3c). These findings establish xCT as necessary for selenium accumulation and selenite toxicity in these models and show that its overexpression is sufficient to confer sensitivity in H1299 cells. Consistent with a requirement for its extracellular substrate, cystine withdrawal reduced selenite sensitivity in A549 cells (Fig. 2f). Thus, selenite toxicity depends on xCT activity and the availability of extracellular cystine.

**Figure 2:**
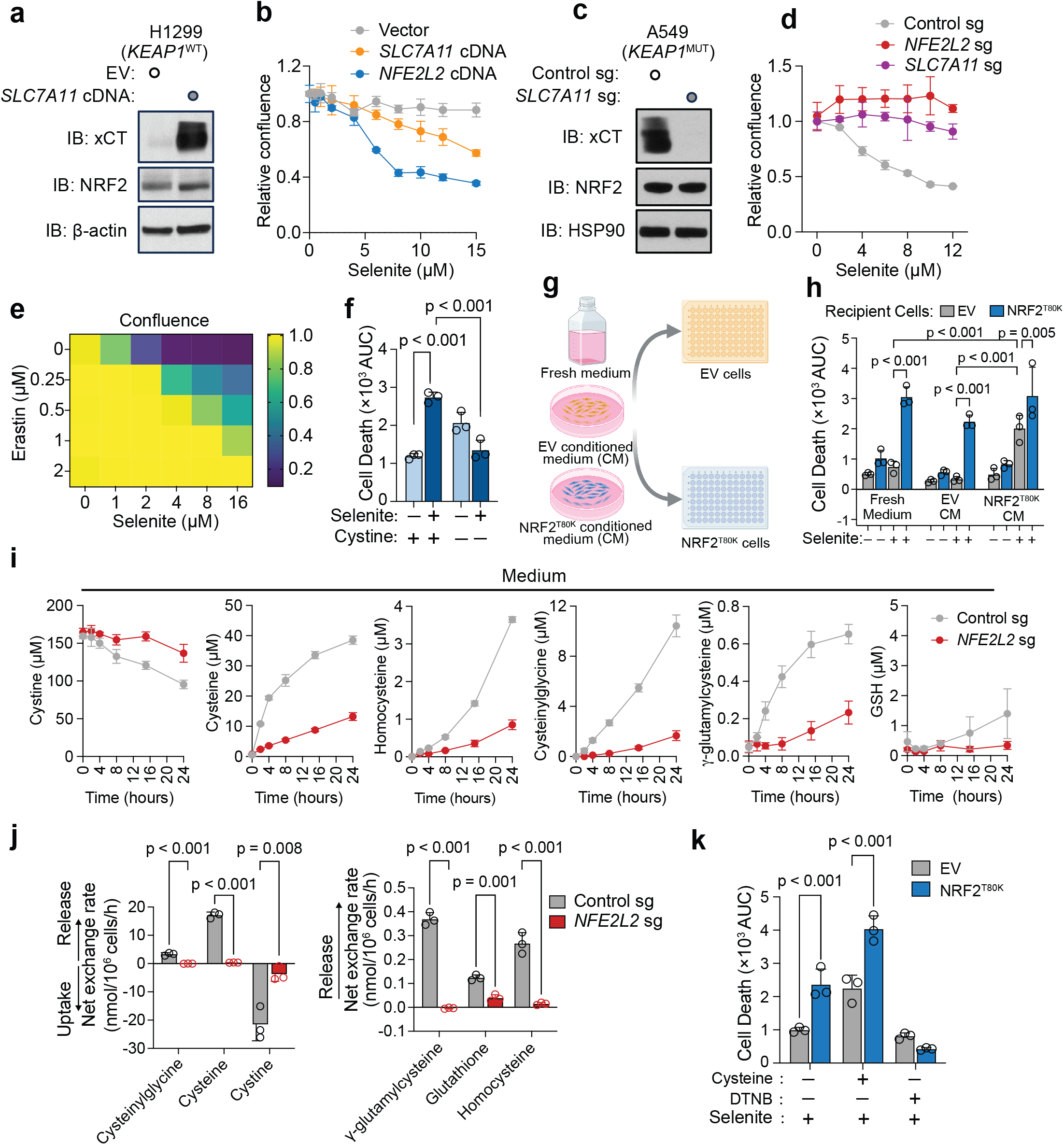
NRF2 drives extracellular reduction of selenite. **a**. Immunoblot of xCT and NRF2 in H1299 cells (*KEAP1* WT) transduced with EV or *SLC7A11* cDNA; β-actin, loading control. **b**. Dose-response of H1299 cells expressing EV, xCT or NRF2^T80K^ to Na_2_SeO_3_, measured as relative confluence by live-cell imaging. **c**. Immunoblot of xCT and NRF2 in A549 cells (*KEAP1*-mutant) transduced with control sgRNA or *SLC7A11* sgRNA; HSP90, loading control. **d**. Dose-response of A549 cells to Na_2_SeO_3._ Cells were transduced with control sgRNA, *NFE2L2* sgRNA or *SLC7A11* sgRNA. **e**. Relative confluence of A549 cells across a matrix of Na_2_SeO_3_ and erastin concentrations; the color bar denotes confluence. **f**. Cell death in A549 cells treated with vehicle or 8 µM Na_2_SeO_3_ in cystine complete vs. cystine free media, cell death quantified as the AUC. **g**. Design of the conditioned-medium transfer experiment. Fresh medium, EV-conditioned medium (CM) or NRF2^T80K^-CM was conditioned for 4 h, then Na_2_SeO_3_ is incorporated and CM is transferred to EV or NRF2^T80K^ recipient cells. **h**. Cell death in recipient cells from the experiment described in **g**. **i**. Extracellular concentrations of cystine, cysteine, homocysteine, cysteinylglycine, Y-glutamylcysteine and GSH over 24 h in medium from A549 cells transduced with control sgRNA or *NFE2L2* sgRNA. **j**. Net exchange rates for the species listed in **i**, calculated from 0-8 h and normalized to cell number. Positive values denote net release; negative values denote net uptake. **k**. Cell death in H1299 EV and NRF2^T80K^ cells treated with vehicle or 8 µM Na_2_SeO_3_ in the presence or absence of cysteine (35 µM) or DTNB (100 µM). Data in **b, d, f, h, i, j, k** are mean ± s.d. Statistical significance was assessed by two-way ANOVA followed by Tukey’s multiple-comparison test.

To determine whether NRF2 promotes selenite toxicity through changes in the extracellular environment, we transferred conditioned medium between genotypes (Fig. 2g). Medium conditioned by NRF2^T80K^-expressing cells and subsequently supplemented with selenite induced cell death in EV recipient cells, whereas medium conditioned by EV cells did not (Fig. 2h). This indicates that NRF2-dependent changes in the extracellular environment were sufficient to sensitize recipient cells to selenite, prompting us to examine whether this effect was associated with increased thiol availability.

Time-course analysis showed that A549 control cells consumed cystine and progressively accumulated cysteine, homocysteine, cysteinylglycine, γ-glutamylcysteine and glutathione (GSH) in the medium. These changes were markedly attenuated in *NFE2L2*-KO cells (Fig. 2i, Supplementary Table 1). Net exchange rates calculated over the 8 h time course confirmed that NRF2 promoted cystine uptake and the net release of the measured thiol species (Fig. 2j, Supplementary Table 2). Untargeted metabolomics identified cysteine, cysteinylglycine and homocysteine, together with the xCT counter-substrate glutamate, among the extracellular metabolites most strongly influenced by NRF2 in A549 cells (Extended Data Fig. 3d, Supplementary Table 2). These metabolites, along with γ-glutamylcysteine, GSH and oxidized glutathione (GSSG), were also elevated in medium conditioned by NRF2^T80K^-expressing H1299 cells (Extended Data Fig. 3e, f, Supplementary Table 3). Together these findings demonstrate that NRF2 activity establishes a thiol-rich extracellular environment.

Finally, we tested whether extracellular reducing capacity is required for selenite toxicity and sufficient to confer sensitivity. Cysteine supplementation enhanced selenite-induced cell death in both EV and NRF2^T80K^-expressing cells, whereas thiol blockade with DTNB or Cu^2+^-mediated thiol oxidation abolished cell death in both genotypes (Fig. 2k and Extended Data Fig. 3h). These effects were accompanied by changes in selenium accumulation: cysteine supplementation eliminated the difference between EV and NRF2^T80K^-expressing cells, whereas DTNB reduced selenium accumulation in both genotypes (Extended Data Fig. 3g). Thus, extracellular thiol availability determines NRF2-dependent differences in cellular selenium accumulation. Sensitization was also observed with the non-thiol reductant tris(2-carboxyethyl)phosphine (TCEP) and with GSH, a tripeptide thiol with limited membrane permeability (Extended Data Fig. 3i, j). Together with the enhancement of cell death by DTT (Fig. 1l), these findings support a model in which NRF2 establishes an extracellular reducing environment that promotes selenite reduction, cellular selenium accumulation and subsequent toxicity.

## Intracellular glutathione promotes selenite-induced thiol oxidation and cell death

Having established the role of extracellular thiols in selenium uptake, we next examined how NRF2 shapes the intracellular thiol pool. *NFE2L2* disruption in A549 cells reduced intracellular cysteine, GSH, γ-glutamylcysteine, homocysteine and GSSG levels without altering cystine abundance (Fig. 3a, Supplementary Table 2). The largest decreases were observed in cysteine and γ-glutamylcysteine, highlighting the impact of NRF2 loss on glutathione precursors. Untargeted metabolomics likewise identified GSH, cysteine and cysteinylglycine among the metabolites most depleted following *NFE2L2* disruption (Extended Data Fig. 4a, Supplementary Table 2). Conversely, NRF2^T80K^-expressing H1299 cells contained elevated levels of homocysteine, γ-glutamylcysteine and cysteinylglycine relative to EV controls (Extended Data Fig. 4b, Supplementary Table 4). These findings demonstrate that NRF2 sustains intracellular thiol availability in addition to promoting extracellular thiol accumulation.

**Figure 3:**
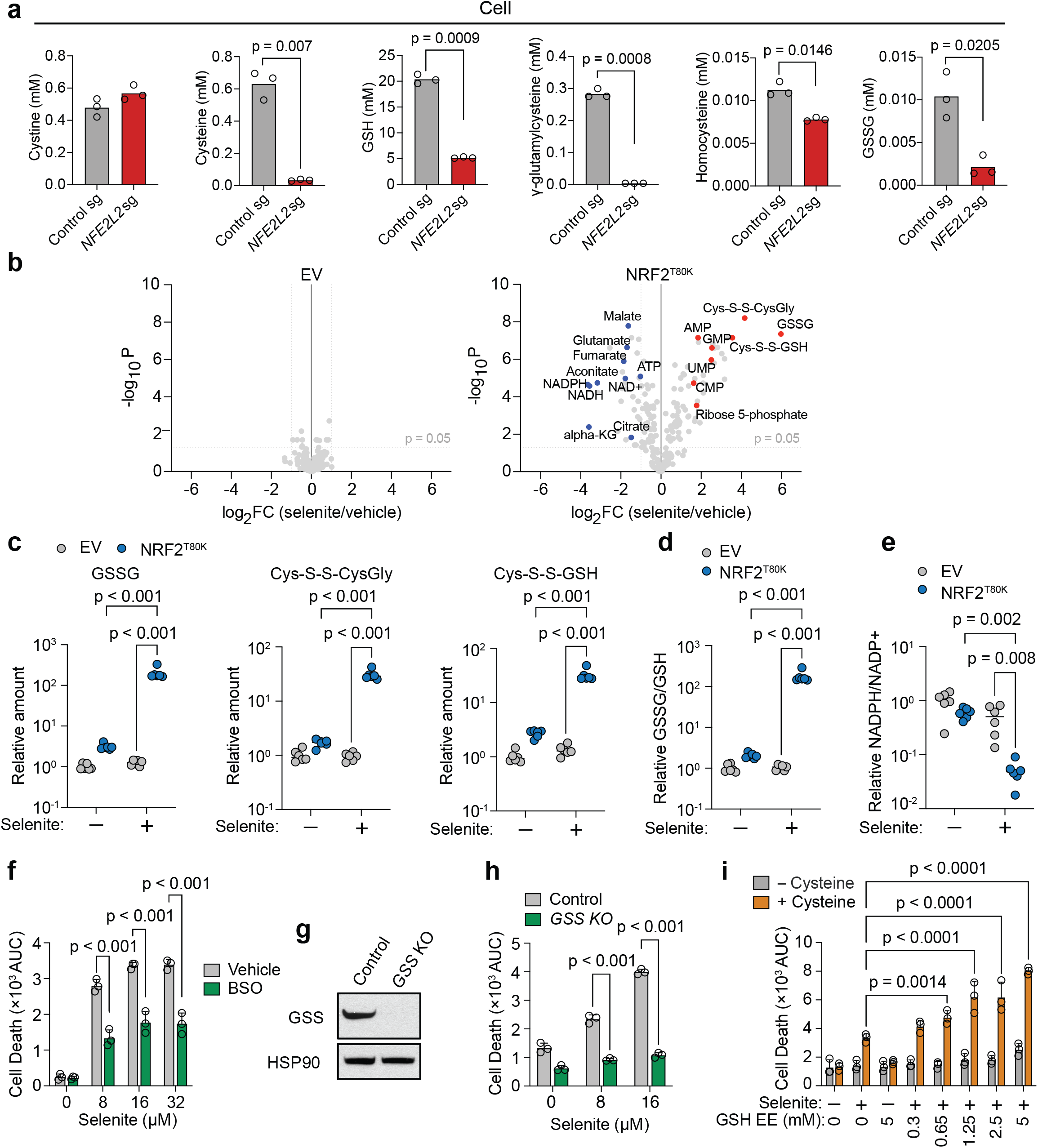
Intracellular glutathione is required for selenite cytotoxicity. **a.** Intracellular concentrations of cystine, cysteine, GSH, Y-glutamylcysteine, homocysteine and GSSG in A549 cells transduced with control sgRNA or *NFE2L2* sgRNA, determined by targeted metabolomics with internal standards. **b.** Volcano plot depicting relative fold change (log_2_) in metabolite abundance versus –log(P values) from H1299 EV (left) or NRF2^T80K^ (right) cells treated with to 8 µM Na_2_SeO_3_ for 4 h compared with vehicle. Significantly increased metabolites are shown in red and significantly decreased metabolites in blue; selected species are labelled. Dashed line, p = 0.05. **c**. Relative amounts of GSSG, Cys-S-S-CysGly and Cys-S-S-GSH in the cells analyzed in **b**. **d**. Relative GSSG/GSH ratios in the cells analyzed in **b**, normalized to the mean GSSG/GSH ratio of vehicle-treated EV samples. normalized to the mean GSSG/GSH ratio of vehicle-treated EV samples. **e**. Relative NADPH/NADP^+^ ratio in the cells analyzed in **b**, normalized to the mean NADPH/NADP^+^ ratio of vehicle-treated EV samples. **f.** Cell death in A549 cells treated with indicated concentrations of Na_2_SeO_3_ in the presence or absence of BSO (100 µM), quantified as the AUC curve. BSO was added 18 hours before the experiment, concurrently treated with 8 µM Na_2_SeO_3_ for 4 hours, and re-supplemented for the next 20 hours after Na_2_SeO_3_ washout. **g**. Immunoblot of GSS in control and *GSS*-knockout A549 cells; HSP90, loading control. **h**. Cell death in the cells depicted in **g.** at the indicated Na_2_SeO_3_ concentrations. **i**. Cell death in H1299 cells pre-treated for 18 h with the indicated concentrations of GSH ethyl ester (GSH-EE). GSH-EE was then removed and the cells exposed to vehicle or 8 µM Na_2_SeO_3_ in medium with or without cysteine (35 µM), added to allow selenium entry; GSH-EE was present only during the pre-treatment. Data in **a, c-f, h, i** are mean ± s.d. Statistical significance was assessed by two-way ANOVA followed by Tukey’s multiple-comparison test. Note the log scale in **c, d** and **e**.

Untargeted metabolomics revealed a pronounced response to selenite in NRF2-hyperactive cells. Principal component analysis separated selenite-treated NRF2^T80K^-expressing cells from all other conditions along PC1, which explained 48.2% of the variance, whereas vehicle- and selenite-treated EV cells clustered together (Extended Data Fig. 4c). Consistent with this pattern, selenite caused minimal changes in individual metabolites in EV cells but extensive metabolic remodeling in NRF2^T80K^-expressing cells (Fig. 3b, Supplementary Table 4). Among the metabolites most elevated were the mixed disulfides Cys-S-S-CysGly and Cys-S-S-GSH, together with GSSG (Fig. 3b, c, Supplementary Table 4). This response indicates widespread oxidation of the intracellular thiol pool. The GSSG/GSH ratio increased by more than two orders of magnitude (Fig. 3d, Supplementary Table 4). Cysteine and GSH, which were elevated at baseline in NRF2^T80K^-expressing cells, declined towards EV levels after selenite exposure whereas cystine increased (Extended Data Fig. 4d-f). The NADPH/NADP⁺ ratio remained unchanged in EV cells but fell sharply in NRF2^T80K^-expressing cells (Fig. 3e, Supplementary Table 4), indicating depletion of cellular reducing capacity following selenite exposure.

To test whether intracellular glutathione is required for selenite-induced cell death, we disrupted its synthesis using pharmacological and genetic approaches. Inhibition of glutamate-cysteine ligase with buthionine sulfoximine (BSO) substantially reduced selenite-induced cell death in A549 cells and NRF2^T80K^-expressing H1299 cells (Fig. 3f, Extended Data Fig. 4g). Likewise, genetic ablation of *GSS*, which encodes glutathione synthetase^49^, significantly reduced cell death following selenite exposure (Fig. 3g, h and Extended Data Fig. 4h). These findings demonstrate that glutathione synthesis is required for selenite-induced cell death.

Conversely, we increased intracellular glutathione by preloading cells with cell-permeable glutathione ethyl ester (GSH-EE), which was removed before selenite exposure to avoid direct extracellular interactions between GSH-EE and selenite. Cysteine was included during selenite treatment to support selenium uptake (Extended Data Fig. 3g). Under these conditions, GSH-EE preloading enhanced selenite-induced cell death in a dose-dependent manner. Without cysteine, when selenium uptake was limited, cell death remained low across the GSH-EE concentrations tested (Fig. 3i). The sensitization observed after GSH-EE removal supports an intracellular contribution of glutathione to selenite toxicity. Together with the protection conferred by inhibiting glutathione synthesis, these findings identify intracellular glutathione as a mediator of selenite-induced cell death.

## Selenite induces mitochondrial dysfunction and oxidative stress in NRF2-hyperactive cells

The metabolic response to selenite extended beyond thiol oxidation. In NRF2^T80K^-expressing cells, ATP declined while AMP, GMP, CMP and UMP accumulated. NADH and NAD^+^ decreased alongside NADPH, and levels of the tricarboxylic acid (TCA) cycle intermediates citrate, aconitate, α-ketoglutarate, fumarate and malate also fell (Fig. 3b, Supplementary Table 4). ATP depletion, nucleotide monophosphate accumulation and reduced TCA cycle metabolite abundance suggested impaired mitochondrial energy metabolism, prompting us to examine mitochondrial function directly.

Mitochondrial membrane potential, assessed by the ratio of tetramethylrhodamine methyl ester (TMRM) to MitoTracker fluorescence, remained unchanged during the first hour of selenite exposure. The potential began to decline at 2 h and was almost completely lost by 4 h in A549 control cells. By contrast, *NFE2L2*-knockout cells were largely protected (Fig. 4a). Selenite also impaired mitochondrial respiration. After a 4 h selenite pulse followed by washout, HCC15 control cells exhibited reduced basal oxygen consumption and a minimal response to FCCP, indicating a persistent loss of spare respiratory capacity. Respiration in *NFE2L2*-knockout cells remained largely unaffected by selenite treatment (Fig. 4b). Similar results were observed in A549 cells (Extended Data Fig. 5a). These functional defects were accompanied by changes in mitochondrial morphology: selenite converted the filamentous mitochondrial network in control cells to a punctate pattern, whereas the network remained intact in *NFE2L2*-knockout cells (Fig. 4c). These changes occurred within the 4 h exposure period, preceding the loss of viability measured after washout and supporting mitochondrial dysfunction as an early event in selenite toxicity.

**Figure 4:**
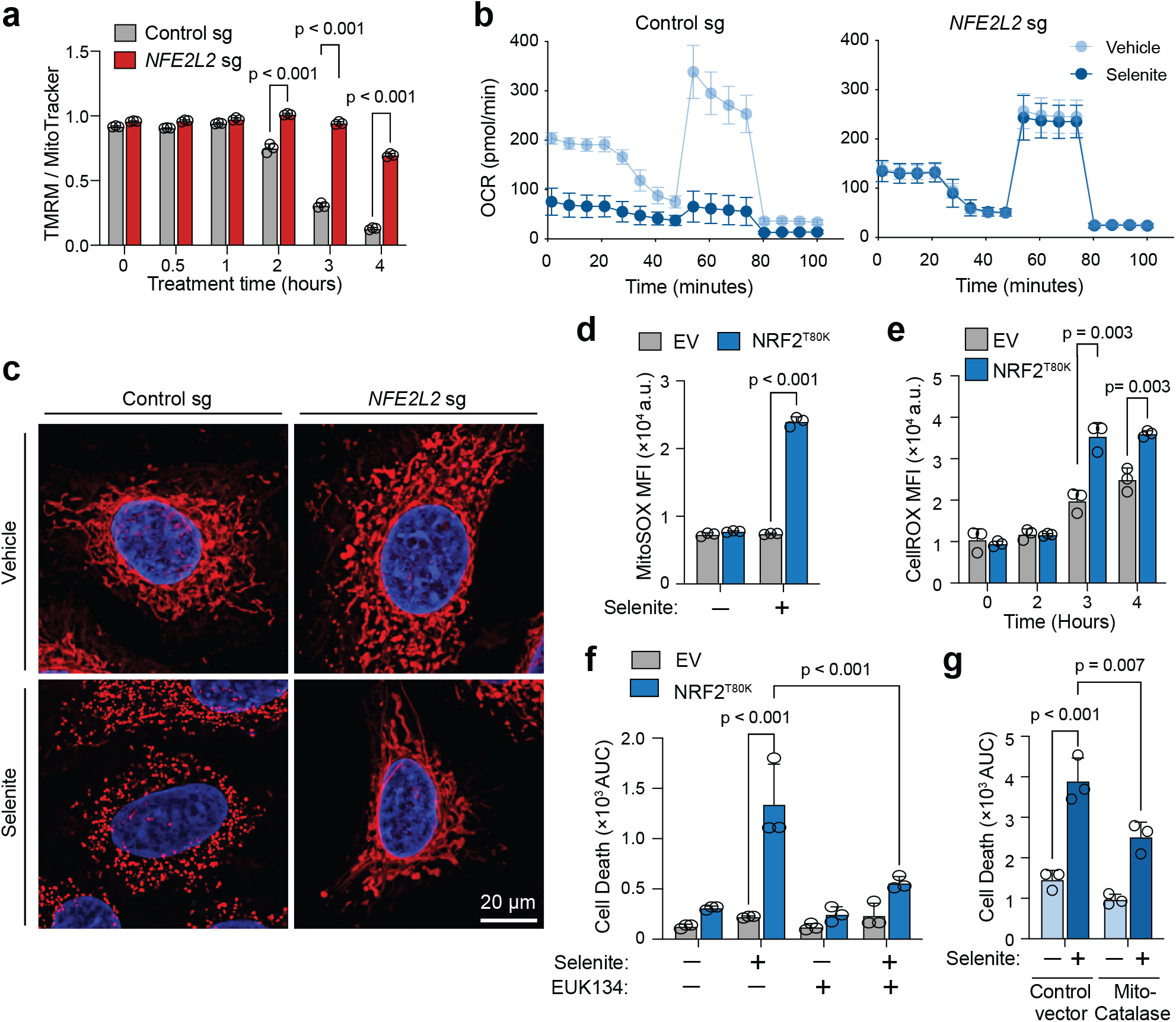
Selenite triggers acute mitochondrial failure and a burst of mitochondrial superoxide. **a.** Mitochondrial membrane potential in A549 cells transduced with control sgRNA or *NFE2L2* sgRNA, measured as the ratio of TMRM to MitoTracker fluorescence, over 4 h of exposure to 8 µM Na_2_SeO_3_. **b.** Oxygen consumption rate (OCR) of HCC15 cells transduced with control sgRNA or *NFE2L2* sgRNA. Cells were treated with 8 µM Na_2_SeO_3_ or vehicle for 4 h, washed, and equilibrated for 60 min in drug-free assay medium before measurement, with sequential injection of oligomycin (1 µM), FCCP (0.5 µM) and rotenone with antimycin A (1 µM each). **c.** Representative confocal images of the cells in **a** stained with MitoTracker (red) and nuclear stain (blue) after 4 h exposure to vehicle or 8 µM Na_2_SeO_3._ Scale bar, 20 µm. **d**. Mitochondrial superoxide in H1299 EV and NRF2^T80K^ cells, measured as MitoSOX median fluorescence intensity, after 4 h exposure to vehicle or 8 µM Na_2_SeO_3._ **e.** Cellular oxidation in the cells described in **d.**measured as CellROX fluorescence intensity, over 4 h of exposure to 8 µM Na_2_SeO_3._ **f.** Cell death in H1299 EV and NRF2^T80K^ cells treated with vehicle or 8 µM Na_2_SeO_3_ for 4 hrs. EUK134 (50 µM) was included in culture media 18 h pre-treatment and 20 h post-treatment to suppress reaction of Na_2_SeO_3_ with free thiols in the media; it was not included during the Na_2_SeO_3_ treatment interval. **g.**Cell death in A549 cells expressing control vector or mitochondrially targeted catalase, treated with vehicle or 8 µM Na_2_SeO_3._ Data are mean ± s.d. Statistical significance was assessed by two-way ANOVA followed by Tukey’s multiple-comparison test. Values in **d** and **e** are median fluorescence intensities from flow cytometry and are compared among conditions.

Mitochondrial dysfunction was accompanied by increased oxidant production. MitoSOX fluorescence, used to assess mitochondrial superoxide, increased following selenite exposure in NRF2^T80K^-expressing H1299 cells but remained unchanged in EV controls (Fig. 4d). CellROX fluorescence was also higher in NRF2^T80K^-expressing cells than in EV controls at 3 and 4 h, indicating increased cellular oxidative stress (Fig. 4e). Hydrogen peroxide levels, assessed using a luminescent probe, likewise increased in selenite-treated NRF2^T80K^-expressing cells (Extended Data Fig. 5b). We tested antioxidant interventions to determine whether oxidative stress contributes to selenite-induced cell death. The superoxide dismutase-catalase mimetic EUK134 markedly reduced selenite-induced cell death in NRF2^T80K^-expressing cells (Fig. 4f). Mitochondrially targeted catalase, which breaks down hydrogen peroxide within mitochondria specifically, also conferred significant protection in A549 cells (Fig. 4g) and NRF2^T80K^-expressing H1299 cells (Extended Data Fig. 5c). These findings support a causal contribution of mitochondrial oxidative stress to selenite-induced cell death.

## Selenite impairs mitochondrial iron-sulfur protein function and abundance in NRF2-hyperactive cells

To further characterize mitochondrial damage in NRF2-hyperactive cells, we profiled the transcriptome after 3 h of selenite exposure, the latest time point at which RNA of sufficient integrity could be obtained. We grouped the NRF2-dependent transcriptional response by subcellular compartment. Non-mitochondrial transcripts remained largely unchanged, whereas nuclear-encoded mitochondrial transcripts declined modestly and the 13 mtDNA-encoded protein-coding transcripts showed a substantially greater decrease (Fig. 5a). All 13 were reduced, with a median ΔΔlog_2_ of approximately –2, compared with –0.18 for nuclear-encoded mitochondrial genes. These findings reveal a preferential loss of mitochondrial genome-encoded transcripts in response to selenite in NRF2-hyperactive cells.

**Figure 5:**
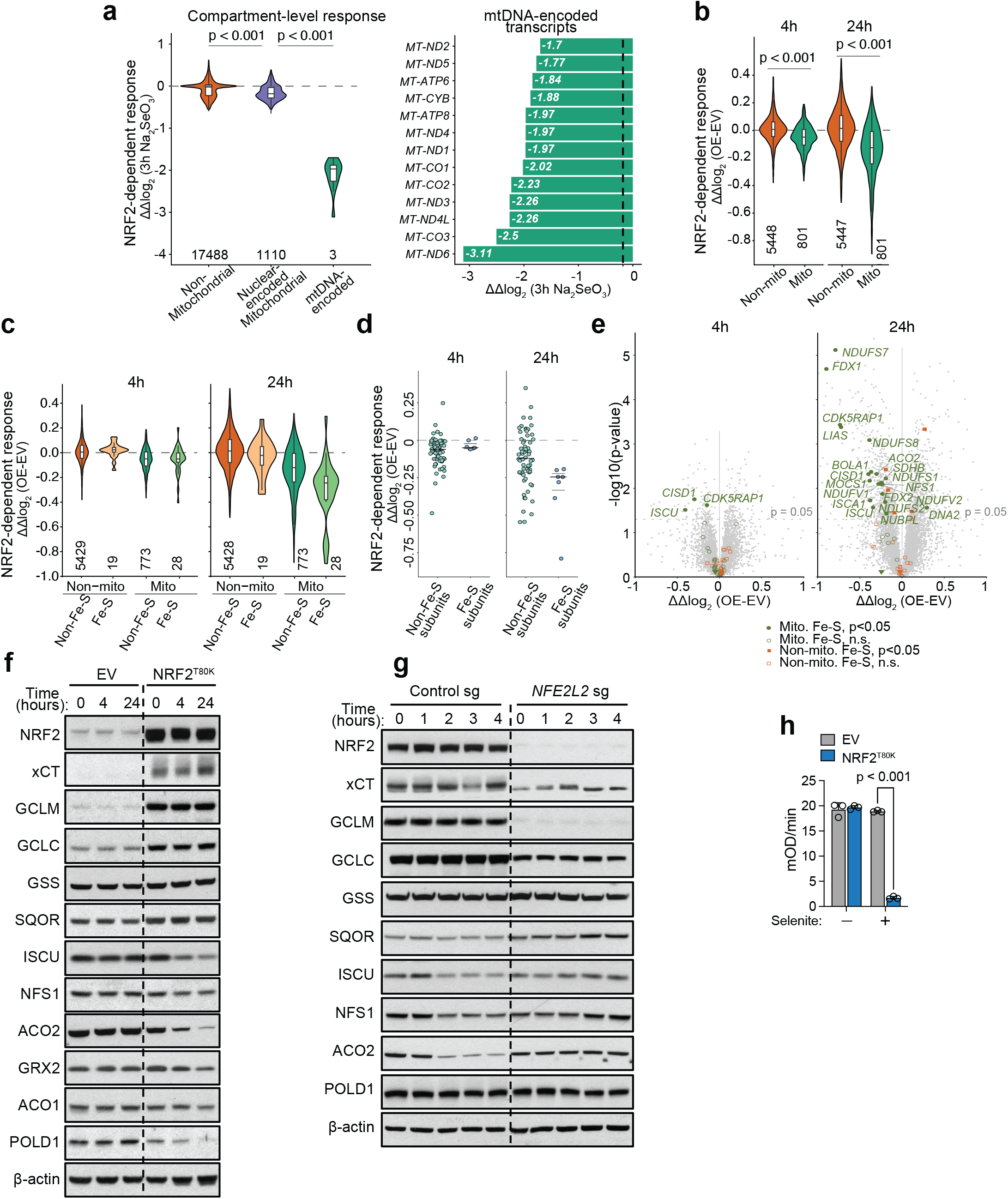
Selenite causes selective loss of iron-sulfur cluster proteins. **a.** Left: NRF2-dependent response of transcript levels (ΔΔlog_2_, [NRF2^T80K^ versus EV] after 3 h, 8 µM Na_2_SeO_3_), grouped as non-mitochondrial (n = 17,488), nuclear-encoded mitochondrial (n = 1,110) and mtDNA-encoded (n = 13). Mitochondrial assignment follows MitoCarta 3.0. Violins are trimmed at the 1^st^ and 99^th^ percentiles; P values, t-test between groups. Right: the same response for each mtDNA-encoded transcript. Dashed line, median of nuclear-encoded mitochondrial genes (–0.18). **b**. NRF2-dependent proteomic response (ΔΔlog_2_, NRF2^T80K^ – EV) for non-mitochondrial (n = 5,448) and mitochondrial (n = 801) proteins, during exposure to 8 µM Na_2_SeO_3_ for 4 h (left) and after 4 h exposure followed by 20 h in drug-free medium (right). P values (Mito versus Non-mito) are shown. **c.** Proteomic data processed as in **b**, with each compartment subdivided into proteins with and without iron-sulfur clusters (non-mitochondrial: n = 5,429 non-Fe-S, 19 Fe-S; mitochondrial: n = 773 non-Fe-S, 28 Fe-S). Violins are trimmed at the top and bottom 1% for groups of 100 or more proteins. **d.** NRF2-dependent response of OXPHOS complex subunits only, separated into non-Fe-S (n = 68) and Fe-S (n = 7) subunits, at the two timepoints. Points are individual subunits; black dashed line, median; grey lines, 25^th^ and 75^th^ percentiles. **e**. Volcano plots of the NRF2-dependent proteomic response (ΔΔlog_2_, NRF2^T80K^ – EV) during exposure to 8 µM Na_2_SeO_3_ for 4 h (left) and after 4 h exposure followed by 20 h in drug-free medium (right). Mitochondrial Fe-S proteins are shown in green and non-mitochondrial Fe-S proteins in orange, filled where P < 0.05 and open where not; all other proteins are grey. Dotted line, P = 0.05. P values are per-protein (pbeta). Gene names are shown for mitochondrial Fe-S proteins with P < 0.05. **f.** Immunoblots of the indicated proteins in H1299 EV and NRF2^T80K^ cells at 0, 4 and 24 h, matching the proteomic time points in **b-e**. Samples were collected before selenite exposure (0 h), at the end of the 4 h exposure (4 h), or after 4 h of exposure followed by 20 h in selenite-free medium (24 h). **g.**Immunoblots of the indicated proteins in A549 cells transduced with control sgRNA or *NFE2L2* sgRNA. Samples were collected at the time points as indicated. **h**. Aconitase activity in H1299 EV and NRF2^T80K^ cells after 4 h exposure to vehicle or 8 µM Na_2_SeO_3,_ measured as the rate of change in absorbance using Aconitase Assay Kit (colorimetric) from Abcam. Data in **h.** are mean ± s.d. Statistical significance was assessed by two-way ANOVA followed by Tukey’s multiple comparison.

We next examined the proteomic response during selenite exposure (4 h) and after washout (4 h exposure followed by 20 h in selenite-free medium, Fig 1i). Mitochondrial proteins showed a greater decline in abundance than non-mitochondrial proteins at both time points, with the difference widening after washout (difference in medians, –0.056 during exposure and –0.141 after washout; Fig. 5b). Thus, mitochondrial protein loss continued after extracellular selenite removal. Within the mitochondrial proteome, iron-sulfur (Fe-S) cluster-containing proteins declined more than other mitochondrial proteins, predominantly after washout. By contrast, non-mitochondrial Fe-S proteins remained unchanged (Fig. 5c)^50^.

Among oxidative phosphorylation (OXPHOS) subunits, the seven Fe-S cluster-containing subunits showed greater depletion than the 68 subunits lacking Fe-S clusters (Fig. 5d, e). This further supports preferential loss of mitochondrial Fe-S proteins. ISCU, the scaffold for nascent [2Fe-2S] cluster assembly, was already reduced during the 4 h selenite exposure, whereas the other Fe-S proteins declined only after washout. This response timeline suggests that disruption of Fe-S cluster biogenesis may precede the broader loss of mitochondrial Fe-S proteins.

We used immunoblotting to track individual proteins over the same time course. In H1299 cells, NRF2, xCT, GCLM and GCLC were more abundant in NRF2^T80K^ expressing cells than in EV cells throughout the time course (Fig. 5f). In NRF2^T80K^-expressing cells, ISCU and ACO2 abundance declined by the end of the 4 h selenite exposure and decreased further during the 20 h after washout, whereas GRX2, a Fe-S protein localized in mitochondria, declined mainly after washout. These decreases were absent in EV cells. NFS1 and SQOR abundance remained relatively stable in H1299 cells, and the extra-mitochondrial Fe-S proteins ACO1 and POLD1 showed no appreciable decline at 4 h. In *KEAP1*-mutant A549 cells, ISCU and ACO2 abundance declined during selenite exposure in control cells and was largely preserved in cells targeted with an *NFE2L2* sgRNA, supporting an NRF2-dependent response (Fig. 5g).

At the end of the 4 h selenite exposure, aconitase activity was nearly abolished in NRF2^T80K^-expressing H1299 cells and remained unchanged in EV cells (Fig. 5h). ACO2 and ACO1 protein were still readily detectable at this time point (Fig. 5f). The near-complete loss of activity while ACO2 protein remained detectable is consistent with damage to the catalytic [4Fe-4S] cluster. Thus, aconitase function was already severely impaired at the end of exposure, before the more pronounced decline in ACO2 abundance after washout.

## SQOR-dependent selenium metabolism links mitochondrial glutathione to selenite toxicity

The preferential disruption of mitochondrial Fe-S proteins prompted us to investigate how selenium is metabolized within mitochondria. Selenide is a close chemical analogue of sulfide, which is oxidized by sulfide-quinone oxidoreductase (SQOR). This enzyme transfers electrons to the coenzyme Q pool and sulfur to an acceptor such as glutathione (Fig. 6a)^51^. SQOR also accepts selenide as a substrate, and the resulting reduction of ubiquinone to ubiquinol has been shown to limit lipid peroxidation and suppress ferroptosis^43^. We therefore asked whether SQOR-mediated selenium metabolism contributes to selenite toxicity in NRF2-hyperactive cells. *SQOR* disruption conferred resistance to sodium selenite in A549 and HCC15 cells (Fig. 6b-d and Extended Data Fig. 6a-c). SQOR loss also reduced selenite-induced mitochondrial superoxide accumulation (Fig. 6e), supporting a role for SQOR upstream of mitochondrial oxidative stress.

**Figure 6:**
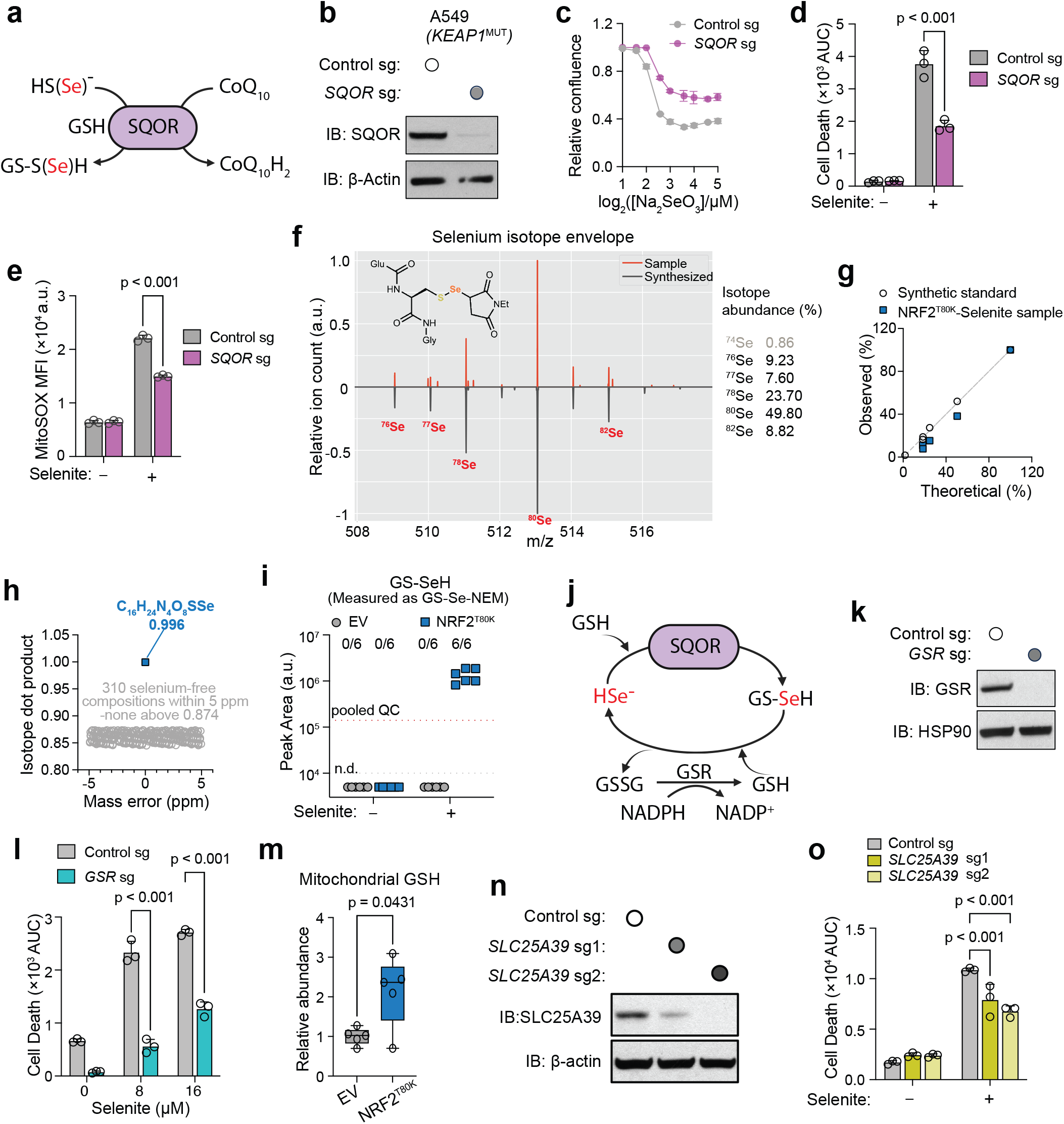
SQOR converts selenide into a reactive glutathione-selenium species. **a**. Schematic illustration for selenide reduction via SQOR: SQOR oxidizes sulfide using coenzyme Q (CoQ) as an electron acceptor and transfers the sulfur atom to glutathione, yielding glutathione persulfide. Selenide enters the same reaction, reducing CoQ and yielding GS-SeH, which is identified in **f-i**. Parenthesized Se denotes the selenium analogue of each sulfur species: HS(Se)^-^, sulfide or selenide; GS-S(Se)H, glutathione persulfide or GS-SeH. **b.** Immunoblot of SQOR in A549 cells transduced with control sgRNA or *SQOR* sgRNA; β-actin, loading control. **c**. Dose-response of the cells in **b** to Na_2_SeO_3_, measured as relative confluence. **d**. Cell death in the cells in **b** after exposure to vehicle or 8 µM Na_2_SeO_3_ quantified as the AUC. **e**. Mitochondrial superoxide in the cells in **b**, measured as MitoSOX median fluorescence intensity, after 4 h exposure to vehicle or 8 µM Na_2_SeO_3_. **f.** Selenium isotope envelope of the NEM adduct of GS-SeH in selenite-treated NRF2^T80K^ cells (sample, upper trace) and in the synthetic standard (lower trace). The table depicts the natural abundance of each selenium isotope. **g.** Observed versus theoretical isotope abundance for the synthetic standard and for the feature detected in selenite-treated NRF2^T80K^ cells. **h**. Isotope dot product of the measured envelope against the predicted formula C_16_H_24_N_4_O_9_SSe (blue) and against 310 selenium-free compositions within 5 ppm of the measured mass (grey). **i**. Peak area of GS-SeH, measured as its NEM adduct, in H1299 EV and NRF2^T80K^ cells exposed to vehicle or 8 µM Na_2_SeO_3_ for 4 h. Fractions above each group give the number of replicates in which the feature was detected. Dashed line, pooled quality-control level; n.d., not detected. **j.** Schematic illustration for proposed Glutathione cycle: SQOR transfers selenium from selenide to glutathione, generating GS-SeH; reaction of GS-SeH with a second glutathione would release selenide and produce GSSG, which GSR reduces back to GSH at the expense of NADPH. Selenium is regenerated rather than consumed at each turn. Selenium-containing species are shown in red. **k.** Immunoblot of GSR in A549 cells transduced with control sgRNA or *GSR* sgRNA; HSP90, loading control. **l**. Cell death in the cells in **k** at the indicated Na_2_SeO_3_ concentrations. **m**. Mitochondrial glutathione levels in untreated H1299 EV and NRF2^T80K^ cells and normalized to median metabolite intensity and expressed relative to EV cells. Box shows the median and quartiles, whiskers the range, and points individual biological replicates. n = 5. **n**. Immunoblot of SLC25A39 in A549 cells transduced with control sgRNA or either of two *SLC25A39* sgRNAs; HSP90, loading control. **o**. Cell death in the cells in **m** after exposure to vehicle or 8 µM Na_2_SeO_3._ Data in **c-e**, **l**, **n** are mean ± s.d. Statistical significance was assessed by two-way ANOVA followed by Tukey’s multiple comparison.; or Welch’s t test.

*SQOR*-mediated electron transfer from selenide to coenzyme Q could increase the ratio of reduced to oxidized coenzyme Q (CoQH_2_/CoQ). However, following selenite exposure, this ratio was lower in NRF2^T80K^-expressing cells than in EV controls (Extended Data Fig. 6d). Antimycin A and rotenone, included as controls, produced the expected shifts in opposite directions. By the 4 h time point, mitochondrial membrane potential and respiration were already severely impaired (Fig. 4a, b). The measured CoQH_2_/CoQ ratio therefore reflects the net redox state of the respiratory chain and cannot distinguish SQOR-mediated electron transfer from the broader effects of mitochondrial dysfunction.

SQOR expression did not correlate with NRF2 status across the NSCLC cell line panel (Extended Data Fig. 6e and Fig 5g), suggesting that NRF2-dependent sensitivity is not driven by increased SQOR abundance. Instead, these findings support a model in which NRF2 promotes cellular selenium uptake while SQOR mediates its subsequent mitochondrial metabolism.

If SQOR transfers selenium to glutathione in a manner analogous to sulfur transfer, the expected product is GS-SeH, a selenium analogue of glutathione persulfide. To test for its formation, we synthesized GS-SeH as an authentic standard and analyzed cellular extracts after derivatizing free thiols and selenols with N-ethylmaleimide (NEM). Extracts from selenite-treated NRF2^T80K^-expressing cells contained a feature with a selenium isotope envelope matching that of the derivatized synthetic standard (Fig. 6f, g). The isotope dot product for the predicted molecular formula C_16_H_24_N_4_O_9_SSe was 0.996; none of the 310 selenium-free candidate compositions within 5 ppm of the measured mass scored above 0.874 (Fig. 6h). This feature was detected in all six replicates of NRF2^T80K^-expressing cells treated with 8 µM sodium selenite for 4 h but was undetectable in EV samples and untreated NRF2^T80K^-expressing cells (Fig. 6i). These findings support the formation of GS-SeH in NRF2-hyperactive cells following selenite exposure.

The formation of GS-SeH provides a potential link between mitochondrial selenium metabolism and glutathione oxidation. Although other thiol-containing metabolites may participate in these reactions, GSH is substantially more abundant than the other measured thiols in NRF2-overexpressing cells, reaching millimolar concentrations. GS-SeH is an established intermediate in glutathione-dependent selenite reduction: selenite reacts with GSH to form selenodiglutathione (GS-Se-SG), which is further reduced to GS-SeH. GS-SeH then reacts with another GSH molecule to produce HSe⁻ and GSSG^36–38^. SQOR-generated GS-SeH could undergo the same reaction, regenerating HSe⁻ that can serve again as a substrate for SQOR. These reactions could therefore establish a redox cycle that repeatedly oxidizes GSH without consuming selenium (Fig. 6j). Glutathione reductase (GSR), present in both the cytosol and mitochondria, replenishes GSH at the expense of NADPH^52,53^, potentially sustaining this cycle after extracellular selenite removal. Consistent with this model, selenite exposure markedly reduced the NADPH/NADP⁺ ratio (Fig. 3e), and *GSR* disruption reduced selenite-induced cell death (Fig. 6k, l).

Because SQOR-mediated selenium metabolism occurs in mitochondria, we next examined glutathione availability within this compartment. Mitochondrial glutathione levels were elevated in NRF2^T80K^-expressing cells relative to EV controls (Fig. 6m, Supplementary Table 5). Mitochondrial GSSG was also detectable in NRF2^T80K^-expressing cells but not in EV cells (Extended Data Fig. 6f, Supplementary Table 5). These findings indicate that NRF2 hyperactivation increases mitochondrial glutathione levels before selenite exposure, providing more substrate for mitochondrial selenium metabolism.

To test whether mitochondrial glutathione availability contributes to selenite toxicity, we targeted its influx into the mitochondria. Mitochondria cannot synthesize glutathione and rely on uptake from the cytosol through SLC25A39 and its paralogue SLC25A40^54^, while GSR maintains the reduced pool by converting GSSG to GSH. Disruption of *SLC25A39* using either of two sgRNAs reduced selenite-induced cell death (Fig. 6n, o). Together with the protection conferred by inhibiting cytosolic glutathione synthesis (Fig. 3f-i), these findings support a model in which cytosol-derived glutathione supplies mitochondrial selenium metabolism. Within mitochondria, SQOR-dependent GS-SeH formation and its subsequent reaction with GSH could sustain selenium redox cycling, supported by GSR-mediated glutathione regeneration, ultimately promoting mitochondrial oxidative stress and cell death.

## Dietary selenite selectively suppresses NRF2-hyperactive tumor growth

To determine whether NRF2-dependent selenite sensitivity extends to tumors *in vivo*, we used the murine KP (*Kras*^LSL-G12D/+^; *Trp53*^fl/fl^) lung tumor cell line, from which the *Keap1*-targeted cells were derived (Extended Data Fig. 1j). Expression of human KEAP1^C273S^, a cysteine mutant that fails to repress NRF2^55^, raised NRF2 protein abundance (Fig. 7a). Mice bearing allografts of KEAP1^C273S^ -expressing or empty-vector control KP cells were maintained on diets containing 0.1 or 2.25 ppm selenium supplied as selenite^56^.

**Figure 7:**
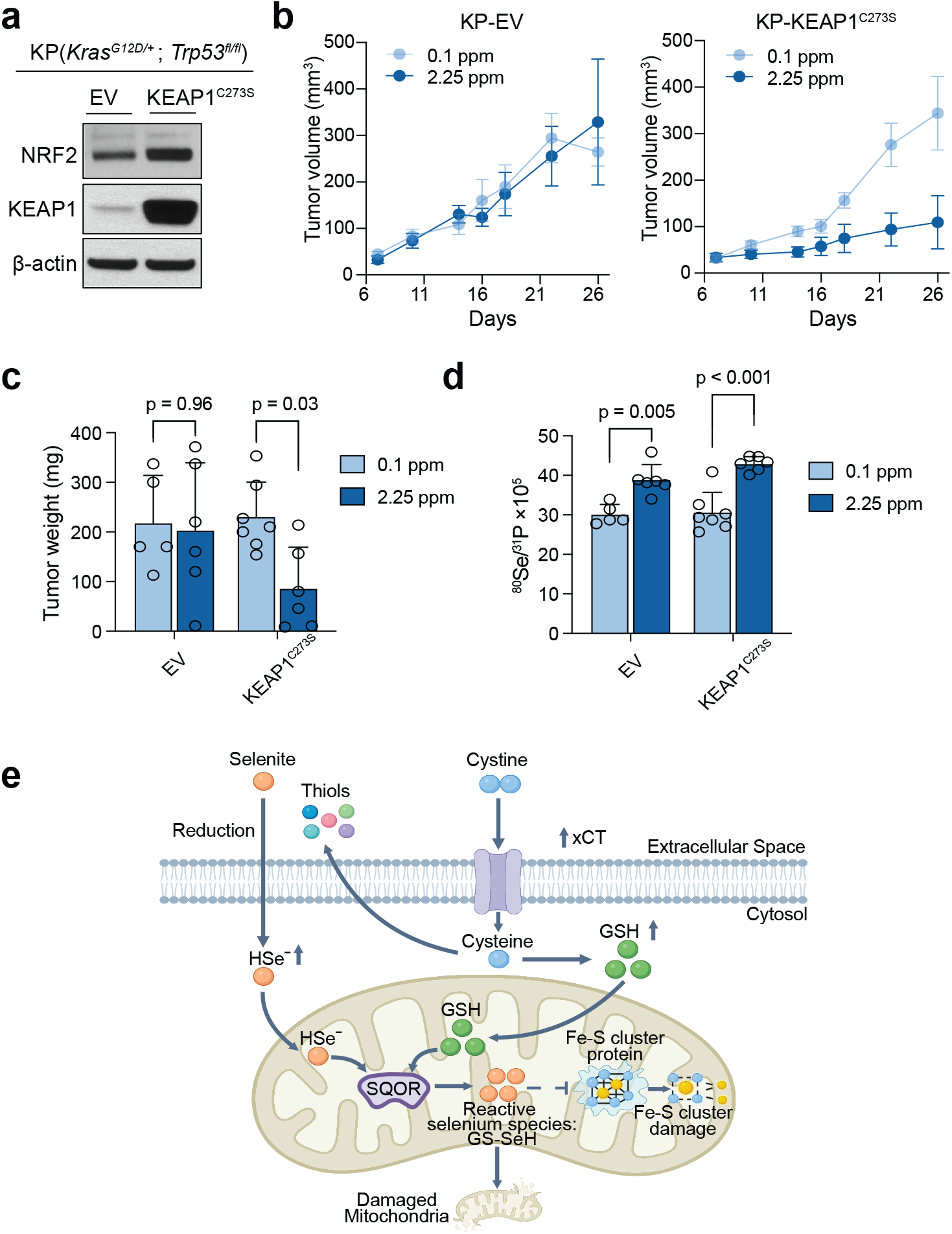
Dietary selenite selectively restrains NRF2-hyperactive tumors. **a.** Immunoblot of NRF2 and KEAP1 in murine KP (*Kras*^G12D/+^; *Trp53*^fl/fl^) lung tumor cells transduced with empty vector or human KEAP1^C273S^ cDNA; β-actin, loading control. This is the same parental line used in Extended Data Fig. 1j. **b**. Growth of subcutaneous allografts of KP-EV (left) and KP-KEAP1^C273S^ (right) cells in mice maintained on diets containing 0.1 or 2.25 ppm selenium. **c.** Final tumor weight for the population described in **b**. **d**. Selenium content of the tumors weighed in **c**, measured by ICP-MS as ^80^Se normalized to ^31^P. **e**. Schematic illustration for the model of NRF2-driven selenite reduction and toxicity: Thiol export reduces extracellular selenite to selenide; selenide enters the cell and reaches mitochondria, where SQOR transfers selenium to glutathione to form GS-SeH; the resulting reactive selenium species destroy iron-sulfur clusters and mitochondrial function fails. Data in b are mean ± s.e.m.; **c-d** are mean ± s.d. Statistical significance was assessed by two-way ANOVA followed by a Tukey’s multiple comparison.

Dietary selenium did not affect the growth or final weight of empty-vector tumors (Fig. 7b,c). In KEAP1^C273S^-expressing tumors, growth trajectories diverged early, and final tumor weight was lower on the selenite-rich diet (Fig. 7b,c). Thus, increased dietary selenite selectively suppressed the growth of NRF2-hyperactive tumors in this model. In tumors that yielded sufficient tissue for endpoint immunoblotting, xCT and GCLM bands were generally stronger in KEAP1^C273S^ tumors than in empty-vector tumors on both diets. GCLC bands also tended to be stronger in KEAP1^C273S^ tumors, particularly in the 0.1 ppm group (Extended Data Fig. 7a). This pattern was consistent with retention of an NRF2-associated thiol metabolism phenotype in vivo. Only two KEAP1^C273S^ tumors from the selenite-rich group yielded sufficient tissue for immunoblotting, so the comparisons involving that group were descriptive.

Tumor selenium content, measured as ^80^Se normalized to ^31^P, increased on the selenite-rich diet in both genotypes (Fig. 7d). These measurements reflect the selenium content of the tissue remaining at endpoint and leave open whether selenium uptake differed between genotypes during tumor growth. No group exhibited body weight loss during the experiment (Extended Data Fig. 7b).

Together, these findings support a model in which NRF2-driven thiol metabolism sensitizes lung cancer cells to selenite (Fig. 7e). NRF2-dependent thiol export promotes extracellular reduction of selenite to selenide, which enters cells and reaches mitochondria. SQOR-mediated transfer of selenium to glutathione generates GS-SeH, which can subsequently react with GSH to regenerate selenide. Coupled to GSR-mediated glutathione recycling, these reactions could sustain a redox cycle that consumes NADPH without consuming selenium. The resulting oxidative stress is associated with rapid mitochondrial dysfunction, impaired Fe-S cluster-dependent enzyme activity and subsequent loss of mitochondrial Fe-S proteins. These effects culminate in cell death that is not prevented by inhibitors of apoptosis, ferroptosis, NLRP3-dependent pyroptosis or RIPK1-dependent necroptosis and lacks the characteristic features of disulfidptosis.

## Discussion

Mutations in *KEAP1* or *NFE2L2* are among the most common genetic alterations in non-small cell lung cancer and define a subset of tumors that remains difficult to treat. Constitutive NRF2 activation increases xCT-mediated cystine import and glutathione synthesis, creating a thiol-rich extracellular environment. Here, we exploit this metabolic state to selectively target NRF2-active cells. These cells reduce selenite extracellularly, take up the resulting selenide, and convert it into GS-SeH through SQOR. The ensuing redox cycle consumes NADPH, damages mitochondrial iron-sulfur clusters, and drives cell death. Thus, the metabolic adaptation that supports resistance to conventional therapies also creates a vulnerability that can be targeted with a compound already administered to patients.

Selenite selectivity for cancer cells has largely been attributed to enhanced uptake. Olm E. *et al*. reported that extracellular reduction by cysteine, cystine import through system xc⁻, and thiol export through multidrug resistance proteins were elevated in chemoresistant cells, concluding that selenium accumulation, rather than intracellular events, determined sensitivity^42^. Our results support the importance of extracellular reduction and identify additional intracellular requirements. Extracellular reduction is required, and xCT is both necessary and largely sufficient to support this process. After uptake, however, glutathione availability and mitochondrial metabolism determine the cytotoxic response: glutathione depletion with BSO or *GSS* KO protects cells, whereas supplementation with GSH-EE before selenite exposure sensitizes them. Disruption of *SQOR*, *GSR*, or *SLC25A39* also reduces cell death. Thus, extracellular chemistry governs selenium delivery, whereas intracellular metabolism determines its consequences. NRF2 supports this pathway at three levels: it promotes the thiol export needed for extracellular selenite reduction, sustains the cytosolic glutathione pool, and increases mitochondrial glutathione, supplying substrate for SQOR-dependent GS-SeH formation. These effects collectively establish a metabolic state that supports both selenium uptake and its conversion into cytotoxic species.

SQOR has recently been shown to protect against ferroptosis by transferring electrons from selenide to ubiquinone, generating ubiquinol that limits lipid peroxidation^43^. Our findings suggest that this protective activity can also exert a cytotoxic effect under conditions of increased selenium flux. SQOR couples ubiquinone reduction to selenium transfer to an acceptor, and the relative consequences of these reactions may depend on substrate availability. At nutritional selenium concentrations, ubiquinol production may favor protection against ferroptosis. In NRF2-hyperactive cells, enhanced extracellular selenite reduction and selenium accumulation may instead drive sufficient GS-SeH formation to promote cell death. Thus, the same enzyme and substrate may produce different outcomes depending on the rate of selenide delivery. We did not detect an increase in the CoQ_10_H_2_/CoQ_10_ ratio after selenite treatment; however, mitochondrial respiration was already impaired at the 4 h time point, complicating interpretation of this result. Moreover, our measurements captured the whole cell CoQ pool. Analysis at earlier time points and within the mitochondrial CoQ pool may better resolve the contribution of SQOR to ubiquinone reduction.

We observed a marked reduction in aconitase activity in NRF2-activated cells following selenite treatment. This functional change provides a more direct indication of the effects of reactive selenium species than changes in protein abundance alone. After 4 h of exposure, ACO2 protein levels remained unchanged, whereas aconitase activity was almost abolished, consistent with damage to its catalytic [4Fe-4S] cluster. This distinction suggests a sequence in which cluster damage causes early loss of enzyme activity, followed by degradation of the resulting apo-proteins over the subsequent 20 h. ISCU was among the earliest Fe-S-associated proteins to decline, with reduced abundance already evident during exposure. As the scaffold for nascent [2Fe-2S] cluster assembly, ISCU contains three cluster-coordinating cysteines: C35, C61 and C104, which are plausible targets of reactive selenium species. Loss of ISCU could further impair cluster biogenesis and replacement, contributing to the delayed, broader depletion of Fe-S proteins. However, the early decline in ISCU could also reflect its turnover kinetics rather than preferential targeting. Distinguishing these possibilities will require direct manipulation of ISCU and assessment of its cluster occupancy and cysteine modifications.

Three features may explain the rapid death of NRF2-activated cells following selenite exposure. First, NRF2-driven thiol export promotes extracellular selenite reduction and selenium accumulation, increasing the intracellular burden of reactive selenium species. Second, selenium is regenerated during redox cycling rather than consumed, potentially sustaining toxicity after extracellular selenite has been removed. Third, damage to a catalytic cofactor can impair enzyme function before protein abundance declines, as illustrated by the near-complete loss of aconitase activity despite the presence of ACO2 and ACO1 protein levels at 4 h. Together, these findings suggest that selenium flux helps determine the outcome: chemistry tolerated at nutritional selenium availability may become lethal when amplified by NRF2-dependent thiol metabolism.

Although our model centers on mitochondrial damage, reactive selenium chemistry may extend beyond this organelle. The cytosol of NRF2-hyperactive cells contains abundant glutathione and cysteine that could react with selenide, and we did detect additional selenium-containing species whose identities remain provisional. Cytosolic chemistry may therefore contribute to the response. Nevertheless, our data supports that the damage was concentrated in mitochondria, and disruption of *SQOR*, *SLC25A39* or *GSR* reduced cell death. These findings place mitochondrial selenium metabolism at the center of the lethal response, while leaving open a contribution from reactions elsewhere in the cell.

Despite its mitochondrial dependence, the ensuing cell death does not readily fit established regulated cell death modalities. Caspase inhibition, iron chelation, ferrostatin-1, liproxstatin-1, MCC950 and necrostatin-1s failed to protect the cells, arguing against canonical apoptosis, ferroptosis, pyroptosis or necroptosis under these conditions. Three observations also distinguish this response from disulfidptosis: thiol-reducing agents enhanced rather than prevented death, glucose remained available throughout treatment, and the F-actin network remained intact. The distinction from iron-driven toxicity is particularly relevant because excess cysteine has recently been shown to mobilize ferritin iron and damage mitochondrial Fe-S proteins through SLC25A28-dependent iron import^57^. Although the mitochondrial damage resembles that observed here, deferoxamine did not protect our cells, whereas disruption of SQOR activity or mitochondrial glutathione availability reduced death. These findings support a distinct initiating mechanism involving mitochondrial selenium metabolism.

In summary, our findings identify a metabolic vulnerability in NRF2-hyperactive cancers and support further investigation of selenite as a therapeutic strategy. Several limitations of our current studies should be acknowledged. Tumor selenium measurements at endpoint confirmed increased selenium content following dietary supplementation but cannot reliably establish differences in delivery between genotypes, because cells with the greatest accumulation may have been preferentially lost. The proposed redox cycle, in which selenium is regenerated rather than consumed, is inferred from established selenium chemistry and supported by the requirement for GSH and GSR, but turnover per selenium atom has not been measured directly. Additionally, the early decline in ISCU identifies it as a candidate target without establishing preferential targeting or a causal role in subsequent Fe-S protein loss. Our *in vivo* evidence remains exploratory: a previously reported dietary selenium concentration of 2.25 ppm supplied as selenite^56^ suppressed tumor growth in a genotype-dependent manner in subcutaneous allografts derived from a single murine cell line. Further studies in autochthonous tumor models, with directly administered selenite and in combination with existing therapies, will be needed to establish the broader relevance of these findings. Defining the dose, schedule and therapeutic window will be essential for translating this NRF2-associated metabolic vulnerability into a treatment.

## METHODS

### Cell culture

Throughout the manuscript, all human NSCLC cell lines and HEK293T cells were acquired as a gift from Gina DeNicola (H1299, H23, A549, HCC15, H460, et al). Cell identities were confirmed by short-tandem repeat profiling; cells were regularly tested to be free of Mycoplasma contamination (MycoAlert Mycoplasma Detection Kit, Lonza and MycoStrip, InvivoGen). Cell lines with stable expression of *Streptococcus pyogenes* Cas9 gene were generated by lentiviral transduction, followed by blasticidin selection as reported previously^58^. NSCLC cancer cells were maintained in RPMI 1640 medium (GIBCO, Life Technologies) with 10% heat-inactivated fetal bovine serum (FBS) (F0926). HEK293T were grown in Dulbecco’s Modified Eagle’s Medium (DMEM) with 10% FBS. Cells were incubated in a humidified incubator at 37°C and 5% CO_2_.

### Western blot analysis

Cell samples were rinsed twice with PBS, then collected from the culture dish into PBS by manual scraping with a plastic cell scraper (Corning, 3010). The suspended cells were pelleted by centrifugation a 5000 x g for 30 seconds. The pellet was resuspended in RIPA lysis buffer containing protease and phosphatase inhibitors (Santa Cruz Biotech, sc-24948) and left on ice for 1 hour, agitated by vortexing every 5 minutes. Samples were then sonicated at high power for 5 minutes, in 30 second pulses (Diagenode, B01020014). Samples were centrifuged at maximum speed for 30 minutes to pellet debris, and the supernatant collected. Protein estimation was performed by DC protein assay (BioRad, 5000111). Samples were equalized to 1 µg/uL protein by dilution with RIPA, and 6X SDS Sample Buffer was incorporated to working 1X concentration (50 mM Tris-HCl, 2% SDS, 9% glycerol, 2% β-mercaptoethanol, 0.02% bromophenol blue). Samples were then heated to 95 C for 5 minutes. SDS-PAGE was performed using Bolt (Thermo Fisher, NWO4120) and Nupage (Thermo Fisher, WBT41220) Bis-tris precast gels. Gels were loaded with 20 µg of protein and run at 120 V for 60 minutes. Transfer was performed in tris-glycine buffer (BioRad, 1610734) containing 10% methanol at 350 mA for 60 minutes. The membrane was blocked in 5% nonfat dry milk dissolved in TBS-T (Santa Cruz, sc-362311) for 60 minutes. Membranes were washed 3 times for 10 minutes with TBS-T. Primary antibodies were diluted with 3% BSA dissolved in TBS-T and incubated with the membrane overnight at 4°C. The following day membranes were rinsed with TBS-T 3 times for 10 minutes. Membranes were incubated with the appropriate HRP-conjugated secondary antibody diluted 1:10,000 in 2% nonfat dry milk in TBS-T, for 1 hour. The membranes were rinsed with TBS-T 3 times for 10 minutes once again. Finally, membranes were treated with ECL (BioRad, 1705061) and developed on radio film.

### Lentivirus Generation

The following plasmids were obtained from Addgene: psPAX2 (Addgene, 12260, a gift from Benjamin Gewurz), pCMV-VSV-G (Addgene, 8454), pLX317-EGFP (Addgene, 115439). pLX317_NFE2L2_T80K and pLX317_NFE2L2, pLGB_KEAP1, pLGB_KEAP1_C273S were gifts from Gina DeNicola. NRF2L2, KEAP1, NFE2L2_T80K and KEAP1_C273S was cloned from these vectors to plenti CMV Hygro (Addgene 17446). Lentiviruses were generated by transfecting HEK293T cells overnight with the expression construct plasmid DNA alongside psPAX2 and pCMV-VSV-G packaging plasmids, diluted in jetPRIME buffer (VWR, 76327-130). The next day, the medium was changed to fresh RPMI 1640 (10% FBS). After 24 and 48 hours, the virus containing medium was collected and filtered with a 0.45 µm syringe filter (Thermo Fisher, 9720514) and was supplemented with 8 µg/mL polybrene (Santa Cruz Biotechnology, sc-134220A) before infection. The sgRNAs used in this study are listed in Supplementary Table 6.

### Lentiviral Infection of NSCLC Cells

Individual sgRNA directed CRISPR/Cas9 knockout was performed as previously described^58^. In brief, Cas9+ NSCLC cells were transduced with lentiviruses expressing the sgRNAs targeting genes of interest. Transduced cells were then selected and expanded in the presence of appropriate selection regent (puromycin or hygromycin). On-target CRISPR effects were validated by immunoblot.

### Dead Cell Measurement with IncuCyte

Cells were plated in clear-flat bottom 96 well plates at a density of 10,000 cells/well (with the exception of H1299 at 8,000 cells/well) in 100 µL final volume of medium. The following day, the medium was change to 100 µL of experimental medium containing 20 nM of Sytox Green (Thermo Fisher, S7020), with or without selenite treatment at the indicated concentration for 4 hours, then exchanged with fresh medium for the next 20 hours. In pharmacological inhibition experiments, the cells were incubated with the drug at the indicated concentration for 18 hours preceding the addition of selenite. The drug concentration was maintained for the 4-hour selenite treatment and for 20 hours after selenite washout.

In cystine starvation experiments, RPMI media free of cysteine, cystine and methionine (MP Biomedicals, 1646454) was supplemented with 100 mM methionine and 10% dialyzed FBS (dFBS, F0392) containing 2 mM glutamine with or without 200 mM cystine as indicated. In conditioned medium exchange experiments, 2.5 x 10^6^ cells were plated in 100 mm dish the day before the experiment. The following day, the medium was changed to 10 mL of fresh medium and incubated for 4 hours to condition the medium. The conditioned medium was filtered through 0.45 µM syringe filter and used to treat experimental cells.

The number of dead cells and cell confluence were measured by the IncuCyte S3 live-cell analysis system (Sartorius, Ann Arbor, MI, USA) in a humidified tissue culture incubator at 37°C with 5% CO_2_. Data were acquired with a 4x or 10x objective lens in phase contrast and green fluorescence (Ex/Em: 460/524 nm, acquisition time: 400 ms) channels. Images were acquired from each well at 3-hour intervals. Image and data processing were performed with IncuCyte 2024A software (Sartorius, Ann Arbor, MI, USA). Dead cell number was normalized to cell confluence [Number of Green Object Count Per mm^2^ / Phase Area Confluence (1/mm^2^/ %)]. The area under the curve (AUC) was calculated as the sum of the dead cell numbers at each time point.

### ICP-MS Analysis of Selenium Content

For preparation of cell line samples for ICP-MS analysis, cells were collected by trypsinization and counted using a Countess Automated Cell Counter (Thermo Fisher, AMQAX2000). The suspended cells were pelleted by centrifugation at 1,000 x g for 1 minute then washed twice with PBS and re-pelleted. Cell pellets were digested overnight in 50 µL of concentrated trace metal grade nitric acid (Fisher, A509P212). The following day samples were diluted 25X with ultrapure water containing 4% hydrogen peroxide. The diluted samples were centrifuged at maximum speed for 30 minutes to pellet debris. The supernatant was collected and stored at room temperature until ICP-MS analysis.

For preparation of animal tissue samples for ICP-MS analysis, animal tissues were collected into cryovials and snap frozen with liquid nitrogen. They were then homogenized by manual pulverization with a Biospec Biopulverizer (Biospec, 5912MS). Approximately 20 mg of pulverized tissue was set aside in a separate pre-weighed tube to be processed for ICP-MS analysis, and the remainder was stored at -80 °C. The tube was weighed again to determine precise sample mass. The tissue was suspended in trace metal grade nitric acid at a volume of 2 µL acid per 1 mg tissue and left to digest overnight. The following day samples were diluted 25x with ultrapure water containing 4% hydrogen peroxide. The diluted samples were centrifuged at maximum speed for 30 minutes to pellet debris. The supernatant was collected and stored at room temperature until ICP-MS analysis.

Selenium content measurement was performed using the iCAP TQ ICP-MS system from Thermo Fisher. All samples were measured using a 100 ppb germanium internal standard. Signal quantification was achieved by generating a standard curve from a 10 ppm selenium stock solution (Inorganic Ventures, CMS-4). Acquisition in the KED mode was performed using helium as the collision gas and targeted ^78^Se. Acquisition in the TQ mode was performed using oxygen as a reaction gas and targeted ^80^Se^16^O. Quantified results were normalized by cell number in the case of cell line samples, and sample mass or phosphorus ^31^P in the case of tissue samples.

### Flow Cytometry Analysis

Cells were plated in 6-well dishes at 350,000 cells/well. The following day, the media was aspirated and cells were treated with 8 µM selenite for the indicated amount of time. Following treatment, MitoSOX Red (Thermo Fisher, M36008), CellROX-DR (Invitrogen #C10422), CellROX-Green (Invitrogen #C10444), MitoTracker Green (Invitrogen #M7514) or TMRM (Invitrogen #T668) were added to the cells at a final concentration of 5 µM for 30 min, with the exception of the TMRM concentration of 200 nM. Following the incubation, cells were washed with PBS, detached with trypsin, pelleted by centrifugation in RPMI containing 10% FBS, washed and resuspended in 400 µL FACS buffer (PBS containing 0.5% BSA, 1 mM EDTA and 25 mM HEPES) for flow cytometry analysis. The samples were analyzed on Accuri C6 Plus Personal flow cytometer (BD biosciences). The data was further analyzed with FlowJo (V11) software (FlowJo).

### RNA sequencing (RNAseq)

Total RNAs were isolated using the RNeasy mini kit following the manufacturer’s manual (Qiagen, 74104). An in-column DNA digestion step was included to remove any residual genomic DNA contamination. To construct indexed libraries, 1 µg of total RNA was used for polyA mRNA-selection using the NEBNext Poly(A) mRNA Magnetic Isolation Module (New England Biolabs), followed by library construction via the NEBNext Ultra RNA Library Prep Kit for Illumina (New England Biolabs). Each experimental treatment was performed in biological triplicate. Libraries were multi-indexed, pooled and sequenced on an Illumina NextSeq 500 sequencer using pair-end 75 bp reads (Illumina).

### Generation of 3XHA-EGFP-OMP25 (HA-Mito) expressing cell lines

HA-Mito-expressing cell lines were generated using a previously described retroviral transduction method^59^. Phoenix-AMPHO packaging cells were cultured in DMEM containing 10% fetal bovine serum (FBS) to approximately 90% confluence. For transfection, 6 μg of pMXs-3XHA-EGFP-OMP25 (Addgene #83356) was mixed with polyethyleneimine (PEI) in jetPRIME buffer and incubated for 20 min at room temperature. After 24 h, the medium was replaced with RPMI 1640 containing 10% FBS, and viral supernatants were collected at 48 and 72 h after transfection. H1299 cells were seeded at 1.4 × 10^5^ cells per well in six-well plates and transduced the following day with undiluted viral supernatants containing 8 μg/mL polybrene. Cells were then selected with 4 μg/mL blasticidin for 7 days.

### Isolation of HA-Tagged Mitochondria for Metabolomic Analysis

Cells engineered to express 3 x HA-EGFP-OMP25 were seeded at 4 x 10^6^ cells per 15-cm dish, with two dishes per biological replicate. Cells were collected 48 h post seeding for metabolomics analysis. The succeeding steps were performed in a 4 °C cold room over ice. 5 mM NEM was added to the KPBS (136 mM KCl, 10 mM KH_2_PO_4_, pH 7.25 in LC/MS grade water) used on the day of the experiment. Cells were washed twice with KPBS and then collected into 2 mL of KPBS by scraping. Cells were pelleted by centrifugation at 1,000 x g for 2 minutes and then resuspended in 1 mL of KPBS. The suspended cells were homogenized in a Dounce homogenizer (Bellco Glass, 1984-10002) for 25 strokes with the low-clearance pestle. The homogenate was centrifuged at 1,000 x g for 2 minutes to pellet nuclei and cellular debris, and the supernatant was collected. The supernatant was incubated with 200 µL of pre-washed anti-HA magnetic beads (Thermo Fisher, 88836) with end-over-end rotation at 4 °C for 5 minutes. Beads were magnetically immobilized and washed twice with KPBS. The beads were then resuspended in 1 mL of KPBS and 250 µL of this suspension was set aside for later protein analysis by western blot. The remaining beads were magnetically immobilized and then resuspended in 80% methanol containing 5 mM NEM. The beads were incubated with end-over-end rotation at 4 °C for 5 minutes. After incubation the suspension was centrifuged at max speed for 10 minutes. The supernatant was collected and immediately proceeded to metabolomic analysis.

### Seahorse analyses of mitochondrial function

Oxygen consumption rate (OCR) and extracellular acidification rate (ECAR) were measured using a Seahorse XFe96 Extracellular Flux Analyzer (Agilent Technologies). A549 and HCC15 cells were seeded at 4 x 10^4^ cells per well in Seahorse XF96 cell culture microplates (cat. no. 103794-100; Agilent Technologies) and cultured overnight in RPMI 1640 medium supplemented with 10% fetal bovine serum (FBS). Cells were washed twice with fresh culture medium and treated with 8 μM sodium selenite (Na_2_SeO_3_; cat. no. ICN19474110; MP Biomedicals) for 4 h. Following treatment, cells were washed twice with Seahorse XF Base Medium (cat. no. 103335-100; Agilent Technologies) supplemented with 10 mM D-glucose (cat. no. D16-500; Fisher Chemical) and 2 mM L-glutamine (cat. no. 25030081; Gibco). Cells were incubated in fresh assay medium at 37°C in a non-CO₂ incubator for 60 min before analysis. OCR and ECAR were measured under basal conditions and following sequential injections of oligomycin (1 μM; cat. no. 495455; Calbiochem), FCCP (0.5 μM; cat. no. HY-100410; MedChemExpress), and rotenone and antimycin A (1 μM each; cat. nos. R8875 and A8674, respectively; Sigma-Aldrich). Four measurement cycles were performed at baseline and after each injection. Mitochondrial respiratory parameters were calculated using Seahorse Wave software (Agilent Technologies).

### F-actin staining and confocal fluorescence imaging

1 x 10^5^ cells were seeded in 35-mm glass-bottom dishes with a 20-mm No. 0 glass-bottom well (cat. no. D35-20-0-N; Cellvis) and cultured overnight in RPMI 1640 medium supplemented with 10% FBS. Cells were treated with 8 μM sodium selenite (Na_2_SeO_3_; cat. no. ICN19474110; MP Biomedicals) for 4 h; control cells received an equivalent volume of vehicle (H_2_O). Following treatment, cells were washed with phosphate-buffered saline (PBS; cat. no. SH30028.02; Cytiva) and fixed with 4% paraformaldehyde in PBS (cat. no. J19943.K2; Thermo Fisher Scientific) for 10 min at room temperature. After washing with PBS, cells were permeabilized with 0.5% Triton X-100 (cat. no. 0694-1L; VWR Life Science) in PBS for 10 min. Cells were then stained with Alexa Fluor Plus 647-conjugated phalloidin (1:1000 dilution; cat. no. A30107; Invitrogen, Thermo Fisher Scientific) for 60 min at room temperature in the dark. After washing with PBS, nuclei were counterstained with Hoechst 33342 (cat. no. H3570; Invitrogen, Thermo Fisher Scientific) for 30 min at room temperature in the dark. Samples were washed with PBS, maintained in PBS to prevent drying, and imaged by confocal fluorescence microscopy. Confocal fluorescence images were acquired using a Nikon AXR Confocal with NSPARC at the Nikon Center of Excellence, H. Lee Moffitt Cancer Center. Images were exported and prepared for presentation using Fiji (ImageJ).

### Metabolomics

Metabolomics was performed as previously described^60^.

#### Internal standard preparation

Thiol (-SH)-containing standards homocysteine, L-cysteine, glutathione, cysteinylglycine, γ-glutamylcysteine – as well as cystine and glutathione disulfide were each prepared as 10 mM stock solutions in 10% Formic acid. 5 mM N-ethylmaleimide-d5 (NEM-d5; CDN isotopes, D-6141) was prepared in 10 mM ammonium formate (pH = 7). For each standard, 100 µL of the 10 mM stock was added to 900 µL of 5 mM NEM-D5 solution and incubated at room temperature for 30 min to yield a 1 mM derivatized internal standard. Internal standards were added to 80% methanol containing 5 mM NEM to the concentrations listed below:

| Metabolite Internal Standard | Formula | Source/Catalog No. | Extraction buffer concentration (μM) |
|---|---|---|---|
| Cystine - D4 | C<sub>6</sub>H<sub>8</sub>D<sub>4</sub>N<sub>2</sub>O<sub>4</sub>S<sub>2</sub> | Cambridge Isotope Laboratories, DLM-9812 | 1 |
| Cysteinylglycine-NEM-D5 | C<sub>11</sub>H<sub>12</sub>D<sub>5</sub>N<sub>3</sub>O<sub>5</sub>S | Prepared in-house | 1 |
| L-Cysteine-NEM-D5 | C<sub>9</sub>H<sub>9</sub>D<sub>5</sub>N<sub>2</sub>O<sub>4</sub>S | Prepared in-house | 0.1 |
| γ-glutamyl cysteine-NEM-D5 | C<sub>14</sub>H<sub>16</sub>D<sub>5</sub>N<sub>3</sub>O<sub>7</sub>S | Prepared in-house | 1 |
| Glutathione-NEM-D5 | C<sub>16</sub>H<sub>19</sub>D<sub>5</sub>N<sub>4</sub>O<sub>8</sub>S | Prepared in-house | 1 |
| L-Homocysteine-NEM-D5 | C<sub>10</sub>H<sub>11</sub>D<sub>5</sub>N<sub>2</sub>O<sub>4</sub>S | Prepared in-house | 0.1 |
| Glutathione disulfide-<sup>13</sup>C<sub>4</sub>,<sup>15</sup>N<sub>2</sub> | C<sub>16</sub><sup>13</sup>C<sub>4</sub>H<sub>32</sub>N<sub>4</sub><sup>15</sup>N<sub>2</sub>O<sub>12</sub>S<sub>2</sub> | Cambridge Isotope Laboratories, CNLM-8782 | 1 |

#### Medium samples preparation

Cells were seeded in 6-well plates in three biological triplicates at a density of 0.35 million/well. 10ul of conditioned medium were extracted at the indicated time points. Samples were diluted in extraction buffer (80% methanol containing 0.5 mmol/L ammonium formate, 5 mmol/L N-ethyl maleimide [NEM], and internal standards, pH= 7) to the following final dilutions 1:20. After addition of extraction buffer, samples were vortexed and incubated for 30 min at 4°C, then centrifuged at 13,000 x g for 10 minutes at 4°C. The supernatant was collected and stored at –80°C until LC-MS analysis.

#### Cell samples preparation

Cells were seeded in 6-well plates in three biological triplicates at a density of 0.35 million/well and extracted at the indicated time points. Media was aspirated and cells were washed with DPBS on wet ice. Ice-cold extraction buffer (400 µL; 80% methanol containing 0.5 mmol/L ammonium formate, 5 mmol/L NEM, and internal standards, pH 7) was then added, and plates were incubated for 30 min at 4°C. Cells were scraped, and the supernatant was collected after centrifugation (13,000 x g, 4°C, 10 minutes) and stored at -80°C until LC-MS analysis.

#### LC/MS analysis and data processing

Metabolite quantification was performed by LC-MS technology as previously described30. In brief, chromatographic separation was performed on a Vanquish UPLC system (Thermo Fisher Scientific) equipped with a Waters Atlantis Premier BEH z-HILIC VanGuard FIT column (2.1 mm ID x 150mm, 2.5 µm particle size; Waters), held at 30°C. A 5 µL sample volume was injected via autosampler. Mobile phase A (10 mmol/L ammonium carbonate and 0.05% ammonium hydroxide in water) and mobile phase B (100% acetonitrile) were used with the following gradient: 0-13 minutes, 80% to 20% of B; 13-15 minutes, 20% of B; 15-20 minutes, 80% of B. Metabolites were detected using a Q Exactive HF (QE-HF) Orbitrap mass spectrometer (Thermo Fisher Scientific) with heated electrospray ionization (H-ESI), scanning m/z 65-950 in both positive and negative modes. The following instrument conditions were maintained: capillary temperature: 320°C, voltage: 3.5 kV, mass resolution: 120,000, and AGC target: 3 x 10^6^. Raw data (.raw files) were analyzed in Skyline35 (version 23.1.0.268). Metabolites were identified by matching exact precursor ion m/z (within a 10ppm window) and retention time against an in-house library of authentic standards. Peak area was used as a read-out of metabolite intensity, normalized to internal standards and corrected for dilution factor to determine concentration.

### Xenograft experiments

C57BL/6J mice were bred and maintained in the animal facility at Moffitt Cancer Center. At 6 weeks of age, mice were assigned to 0.1 ppm selenium (cat. no. TD.96363; Teklad, Inotiv) or 2.25 ppm selenium (cat. no. TD.250706; Teklad, Inotiv). Selenium was supplied in the form of sodium selenite. After 5 weeks on the assigned diet, mice were inoculated subcutaneously in the lower dorsal flank near a hind limb with 2.5 x 10^5^ KP1.9 cells expressing either empty vector (EV) or KEAP1-C273S, suspended in 100 μL PBS. Mice continued receiving their assigned diets after tumor implantation. Beginning 7 days after implantation, body weight and tumor dimensions were recorded three times per week. Tumor volume was calculated as (length × width^2^)/2. Mice were euthanized at the experimental endpoint following the development of tumor ulceration. Tumors were excised, weighed, and photographed. Tumor tissues were pulverized on dry ice before selenium was quantified by inductively coupled plasma mass spectrometry (ICP-MS).

Other materials and reagents used in this study are listed in Supplementary Table 7.

### Statistical analysis

Unless otherwise indicated, all bar graphs and line graphs represent the arithmetic mean of at least three independent experiments (n = 3), with error bars denoting standard deviations. Data were analyzed using two-tailed paired Student t test or analysis of variance (ANOVA) with the appropriate post-test using GraphPad Prism7 software.

## Supporting information

Materials and Reagents

sgRNA primer list

Mito-IP Metabolomics

H1299_selenite_metabolomics

H1299_metabolomics

A549_8h_metabolomics

A549_time course_medium_metabolomics

## Graphics

Figures were drawn with GraphPad, Biorender and Adobe Illustrator.

## Supplementary Tables

## Distribution of Materials

Correspondence and material requests should be addressed to.

## Data availability

The RNAseq datasets have been deposited to the NIH GEO omnibus and will be released upon publication. The proteomics datasets have been deposited to the PRIDE and will be released upon publication.

## Acknowledgements

We thank Gina M. DeNicola for helpful suggestions and advice on thiol analysis and shared reagents; John Cleveland and Florian Karreth for editing and proofreading; Marcelo Bonini for advice on the use of ROS scavengers; Ana Gomes for sharing cell line and reagent; Nathan Ward for providing sgRNAs targeting *SLC25A39*; and Elsa Flores for supporting seahorse analysis; Dalia Ercan for proteomic data analysis. This work was supported by the National Cancer Institute (R00CA270423 to C.J.), and National Institute of General Medical Sciences (R35GM165628) of the National Institutes of Health. We acknowledge the use of Chatgpt (GPT-6.1 Sol-Light) for grammar checking and editing. This work was also supported by the Moffitt Microscopy, Proteomics/Metabolomics, Genomic Cores, Biostatistics and Bioinformatics Shared Resource (BBSR), which are funded in part by Moffitt’s Cancer Center Support Grant (NCI, P30CA076292).

## Contributions

C.J. and T.C. conceived the study, designed the experiments, and interpreted the results. T.C. performed cell culture, western blotting, cell death and ROS assays, metabolomics, RNA sequencing, proteomics, and molecular cloning. R.M. performed mitochondrial isolation, western blotting, and ICP-MS analyses. Y.H. performed mitochondrial isolation, cell proliferation, and in vivo experiments. J.L. performed western blotting, generated cell lines, and managed mouse colonies. N.T. and H.S. synthesized selenium-containing metabolites, generated standards, and performed metabolomics analyses. Y.T., S.A., and M.T. performed bioinformatics analyses. B.A.Y. and F.K.C. assisted with metabolomics analyses. S.P. works on graphic figures and supports molecular assays. R.P. supports in vivo experiment, and R.P. and A.P. extracted tissue samples for ICP-MS analyses. M.L., B.F., and J.K. supported metabolomics and proteomics analyses. J.S. and W.G.S. supported confocal imaging analyses. C.J. wrote the first draft with input from T.C., R.W, Y.H and J.L. All authors read and approved the manuscript. C.J. supervised the study and secured funding.

## Ethics declarations

**Extended Data Figure 1:**
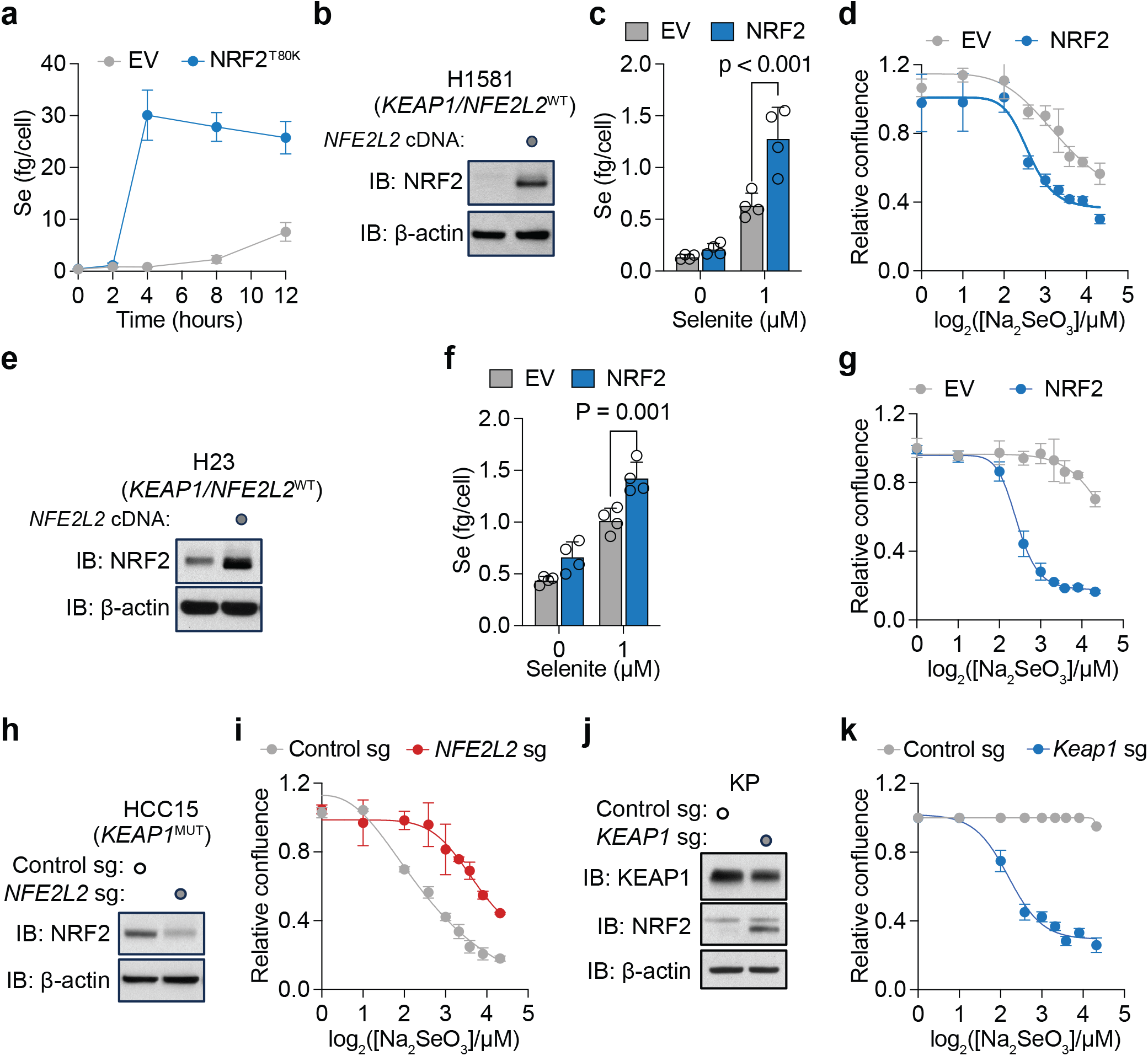
NRF2 promotes selenium accumulation and selenite sensitivity across NSCLC genetic backgrounds. **a**. Time-course of intracellular selenium in H1299 cells expressing EV or NRF2^T80K^, measured by ICP-MS via ^78^Se, following exposure to 4 µM Na_2_SeO_3._ **b**. Immunoblot of NRF2 in H1581 cells transduced with EV or WT human *NFE2L2* cDNA; β-actin, loading control. **c**. Intracellular selenium levels of cells in **b** after 4 h exposure to the indicated Na_2_SeO_3_ concentrations. **d**. Dose-response of cells in **b** to Na_2_SeO_3_, measured as relative confluence. **e-g.** Analogous data to **b-d** for H23 cells. **h**. Immunoblot of NRF2 in HCC15 cells transduced with either control sgRNA or *NFE2L2* sgRNA. **i**. Dose-response of cells in **h** to Na_2_SeO_3_. **j**. Immunoblot of KEAP1 and NRF2 in a murine KP (*Kras*^LSL-G12D/+^; *Trp53*^fl/fl^) lung tumor line transduced with control sgRNA or *Keap1* sgRNA. **k**. Dose-response of the cells in **j** to Na_2_SeO_3_. Data in **a, c, d, f, g, i, k** are mean ± s.d. Statistical significance was assessed by two-way ANOVA followed by a Tukey’s multiple comparison.

**Extended Data Figure 2:**
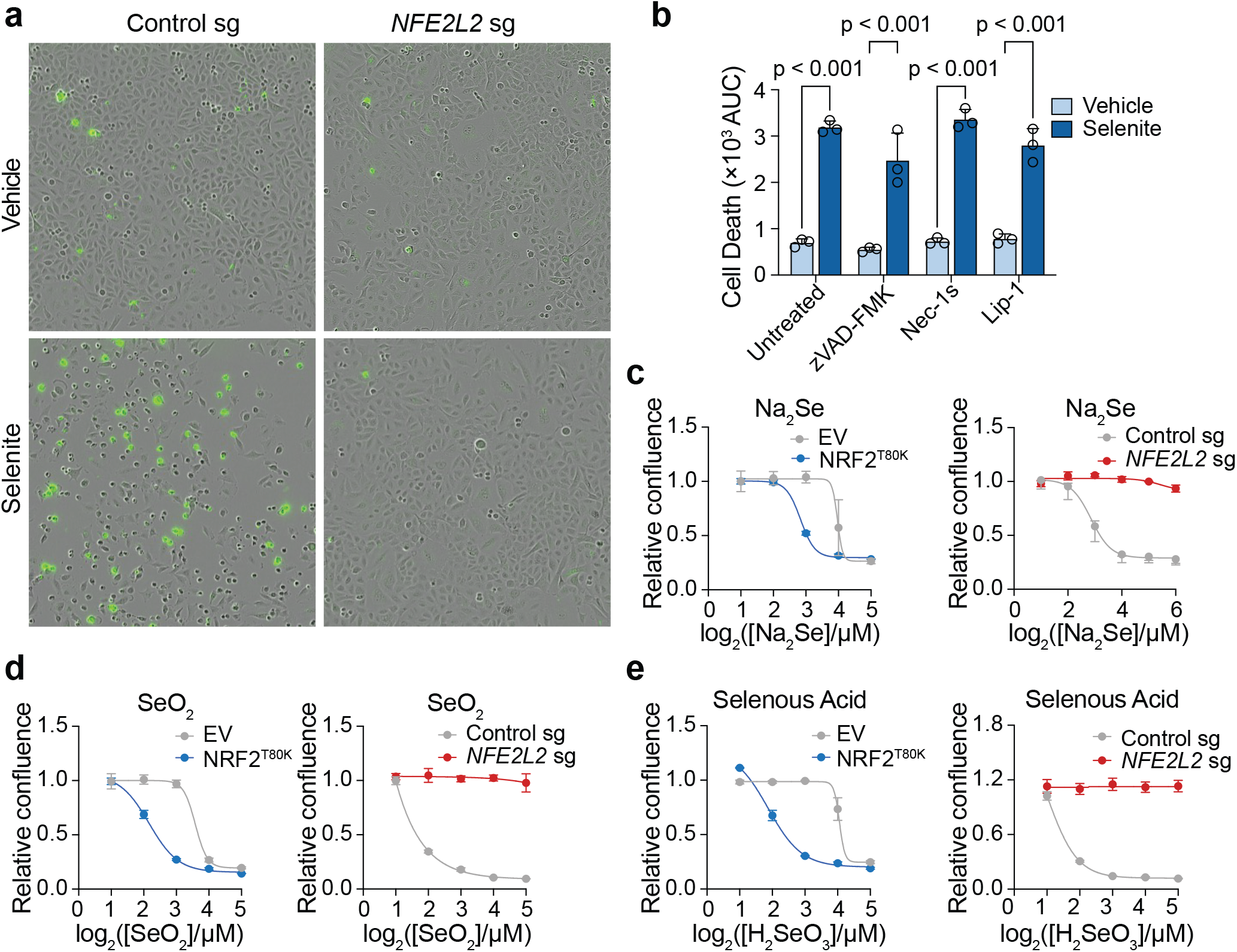
Selenite-induced death is independent of canonical death pathways and extends to other inorganic selenium species. **a.** Representative phase-contrast images from IncuCyte S3 overlaid with SYTOX Green fluorescence in A549 cells transduced with control sg or *NFE2L2* sgRNA, 20 h after a 4 h exposure to the vehicle or 8 µM Na_2_SeO_3_. **b.** Cell death in A549 cells, quantified as AUC of SYTOX Green positive cell counts normalized to total cell area over time, following a 4 h exposure to vehicle or 8 µM Na_2_SeO_3_ in the presence of zVAD-FMK (25 µM), necrostatin-1s (25 µM) or liproxstatin-1 (2.5 µM). **c-e.** Dose-response of H1299 EV and NRF2^T80K^ cells (left) and A549 control sgRNA and *NFE2L2* sgRNA cells (right) to Na_2_Se (**c**), SeO_2_ (**d**) and selenous acid (H_2_SeO_3_) (**e**), measured as relative confluence. Data in **b-e.** are mean ± s.d. Statistical significance was assessed by two-way ANOVA followed by a Tukey’s multiple comparison.

**Extended Data Figure 3:**
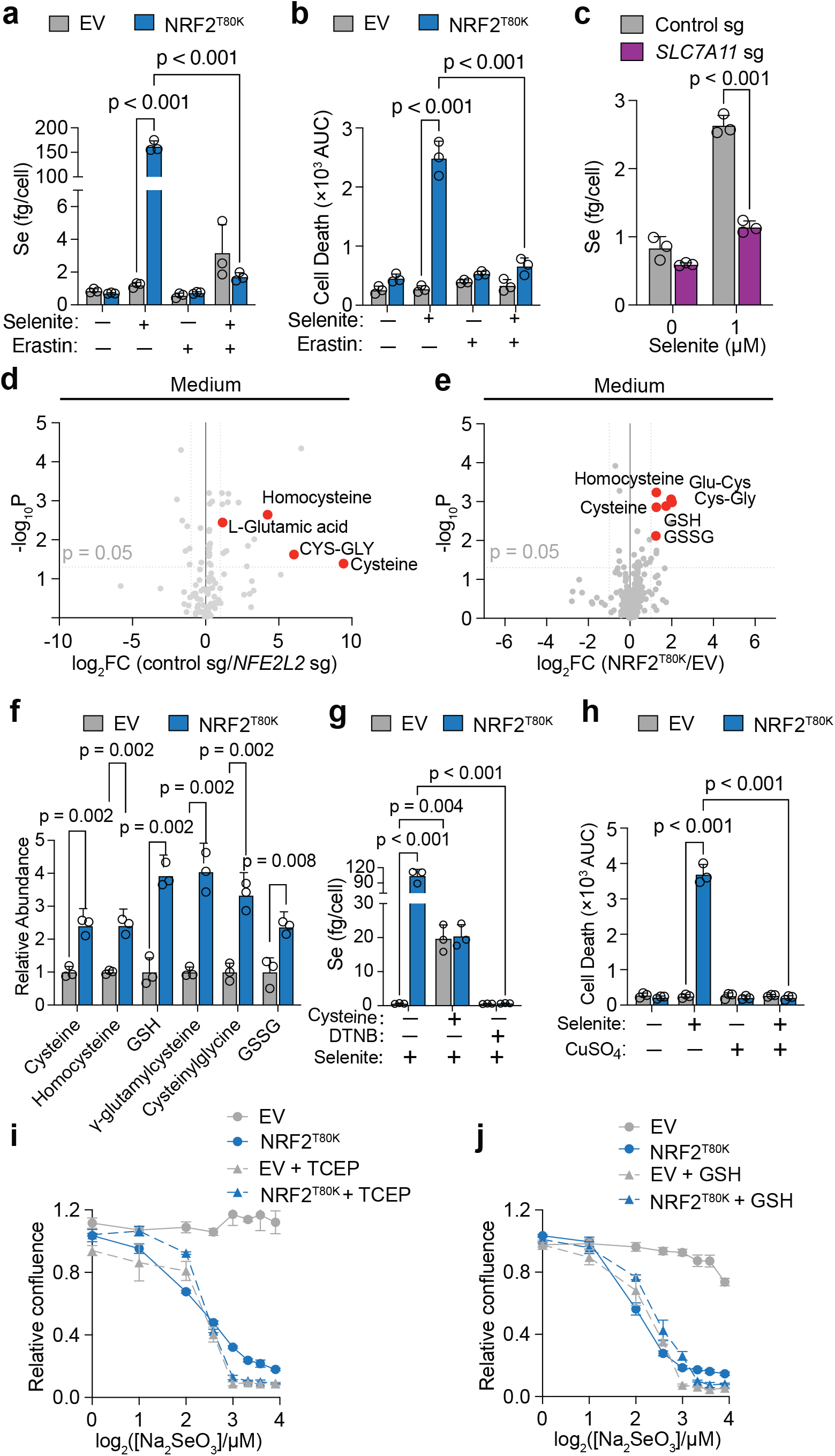
xCT-dependent thiol export determines selenium delivery. **a.** Intracellular selenium in H1299 EV and NRF2^T80K^ cells treated with vehicle or 8 µM Na_2_SeO_3_ in the presence or absence of erastin (5 µM). **b**. Cell death quantification of the cells depicted in **a**. **c**. Intracellular selenium in A549 cells transduced with control sgRNA or *SLC7A11* sgRNA after 4 h exposure to the indicated Na_2_SeO_3_ concentrations, measured by ICP-MS via ^78^Se. **d**. Volcano plot of extracellular metabolites in medium conditioned by A549 cells transduced with control sgRNA or *NFE2L2* sgRNA; thiol species and glutamate are highlighted. Dashed line, p = 0.05. FC: fold change. **e**. Volcano plot of extracellular metabolites in medium conditioned by H1299 EV versus NRF2^T80K^ cells; thiol species are highlighted. Dashed line, p = 0.05. FC: fold change. **f**. Relative abundance of thiol species in medium conditioned by H1299 EV and NRF2^T80K^ cells. **g**. Intracellular selenium in H1299 EV and NRF2^T80K^ cells after exposure to 8 µM Na_2_SeO_3_ in the presence or absence of cysteine (35 µM) or DTNB (100 µM). **h**. Cell death in H1299 EV and NRF2^T80K^ cells treated with vehicle or 8 µM Na_2_SeO_3_ in the presence or absence of CuSO_4_ (15 µM). **i**. Dose-response of H1299 EV and NRF2^T80K^ cells to Na_2_SeO_3_ with or without TCEP (100 µM). **j**. Cells treated as in **i** with extracellular GSH (1mM). Data in **a-c**, **f-j** are mean ± s.d. Statistical significance was assessed by two-way ANOVA followed by a Tukey’s multiple comparison.

**Extended Data Figure 4:**
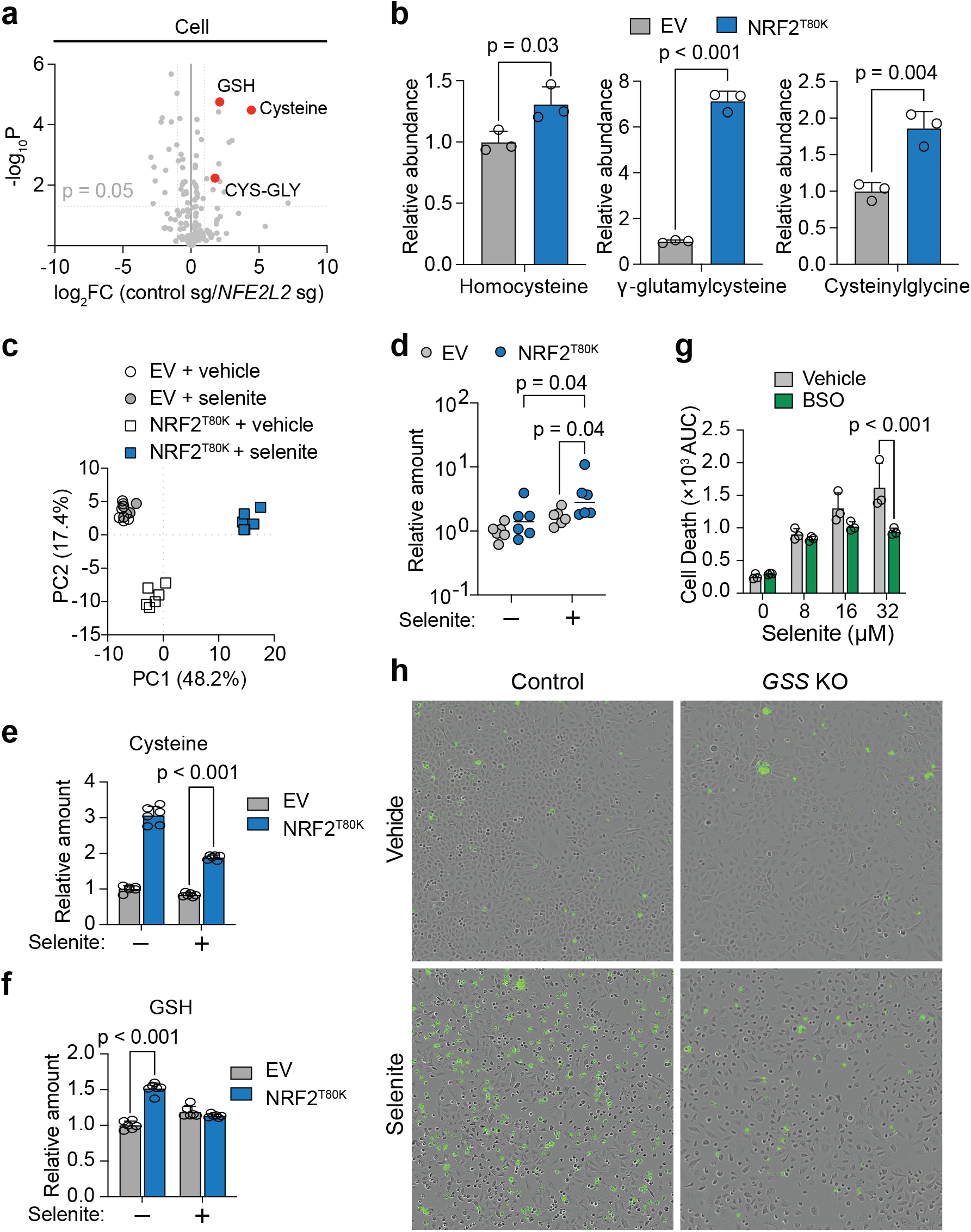
NRF2 sustains the intracellular thiol pool. **a.** Volcano plot of intracellular metabolites in A549 control sgRNA versus *NFE2L2* sgRNA cells; GSH, cysteine and cysteinylglycine are highlighted. Dashed line, p = 0.05. **b.** Relative abundance of homocysteine, Y-glutamylcysteine and cysteinylglycine in H1299 EV and NRF2^T80K^ cells. **c**. Principal component analysis of the intracellular metabolomes of H1299 EV and NRF2^T80K^ cells treated with vehicle or 8 µM Na_2_SeO_3_; PC1 and PC2 account for 48.2% and 17.4% of the variance, respectively. **d-f**. Relative levels of intracellular cystine (**d)**, cysteine (**e**), and GSH (**f)** of the cells depicted in **c**. **g**. Cell death in H1299 NRF2^T80K^ cells treated with indicated concentrations of Na_2_SeO_3_ in the presence or absence of BSO (100 µM), quantified as by AUC. **h.** Representative phase-contrast images overlaid with SYTOX Green fluorescence in control and *GSS*-knockout cells after exposure to vehicle or 8 µM Na_2_SeO_3_. Data in **b**, **d-g** are mean ± s.d. Statistical significance was assessed by two-way ANOVA followed by a Tukey’s multiple comparison.

**Extended Data Figure 5:**
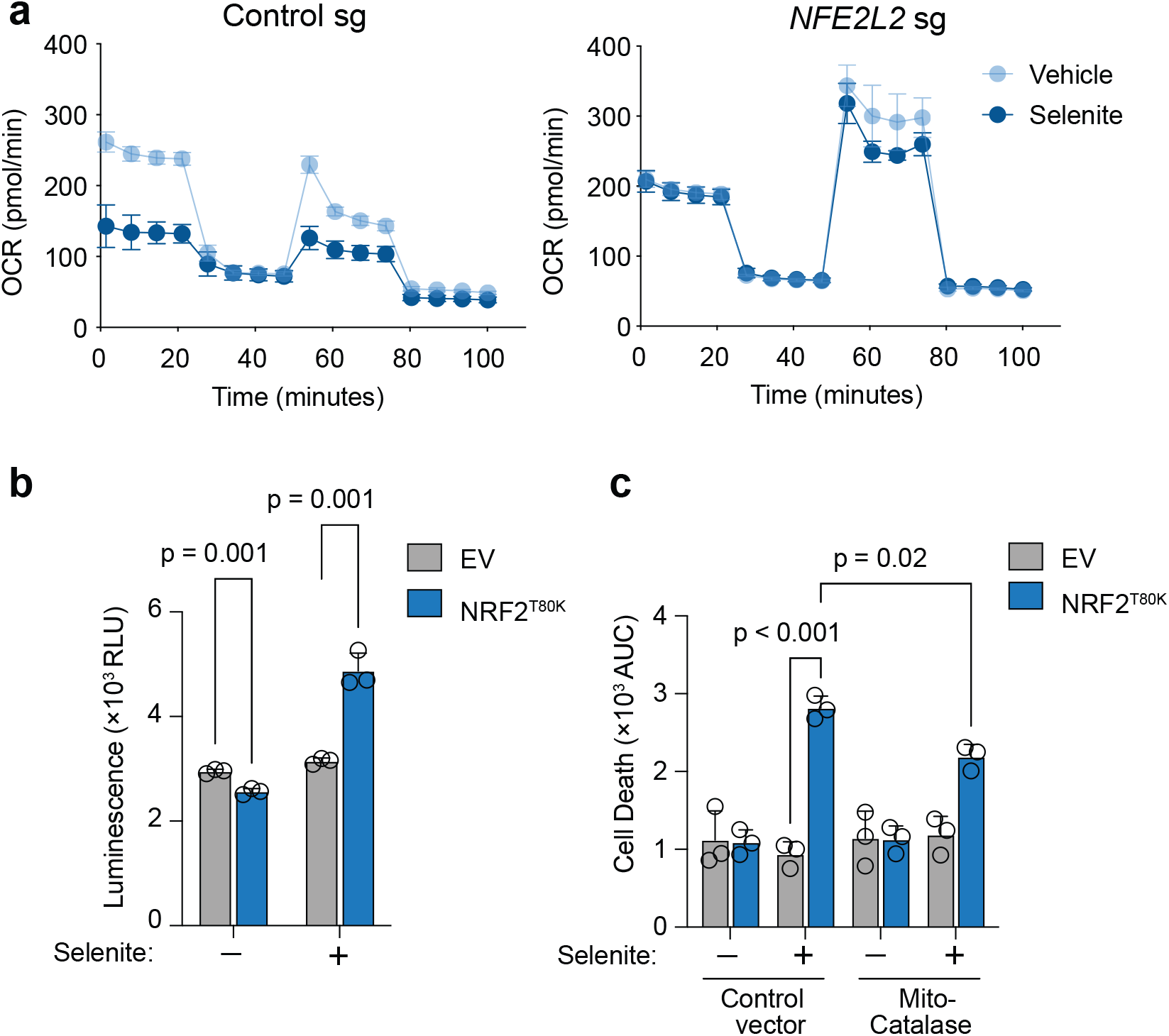
Mitochondrial respiration and oxidant production after selenite exposure. **a.** Oxygen consumption rate of A549 cells transduced with control sgRNA or *NFE2L2* sgRNA, treated and measured as in Fig. 4b. **b**. Hydrogen peroxide in H1299 EV and NRF2^T80K^ cells after 4 h exposure to vehicle or 8 µM Na_2_SeO_3_ measured with Promega ROS-Glo™ H_2_O_2_ and expressed as luminescence units. **c.** Cell death in H1299 cells expressing control vector or mitochondrially targeted catalase, treated with vehicle or 8 µM Na_2_SeO_3_. Data are mean ± s.d. Statistical significance was assessed by two-way ANOVA followed by a Tukey’s multiple comparison.

**Extended Data Figure 6:**
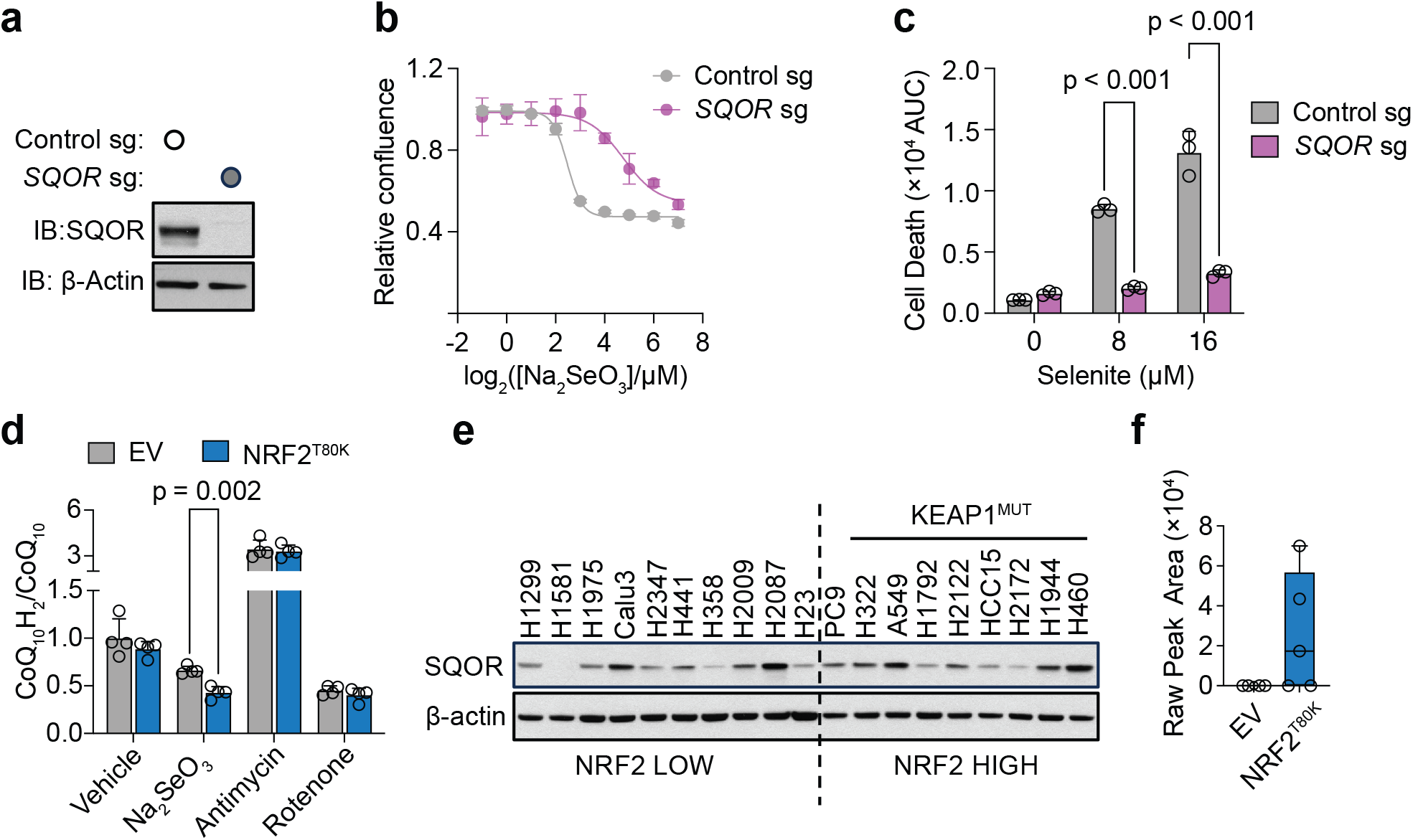
SQOR is required for selenite toxicity and is not an NRF2 target. **a**. Immunoblot of SQOR in HCC15 cells transduced with control sgRNA or *SQOR* sgRNA; β-actin, loading control. **b.** Dose-response of the cells depicted in **a** to Na_2_SeO_3_, measured as relative confluence. **c.** Cell death in the cells depicted in **a.** at the indicated Na_2_SeO_3_ concentrations, quantified as the AUC. Statistical significance was assessed by two-way ANOVA followed by a Tukey’s multiple comparison. **d**. Ratio of reduced to oxidized coenzyme Q (CoQ_10_H_2_/CoQ_10_) in H1299 EV and NRF2^T80K^ cells after 4 h exposure to vehicle or 8 µM Na_2_SeO_3_. Antimycin A (1 µM) and rotenone (1 µM) are included as assay controls, confirming assay sensitivity to both an increase and a decrease in the ratio. Statistical significance was assessed by multiple t-test. **e**. Immunoblot of SQOR across a panel of NSCLC cell lines grouped by NRF2 status; *KEAP1*-mutant lines are indicated. β-actin, loading control. **f**. Mitochondrial GSSG in untreated H1299 EV and NRF2^T80K^ cells, shown as raw peak area because the species was below the limit of detection in EV samples. Boxes show the median and quartiles, whiskers the range, and points individual biological replicates; 3 of 5 NRF2^T80K^ replicates were above the limit of detection. Data in **b**, **c** and **d** are mean ± s.d. Statistical significance in **c**. was assessed by two-way ANOVA followed by a Tukey’s multiple comparison.

**Extended Data Figure 7:**
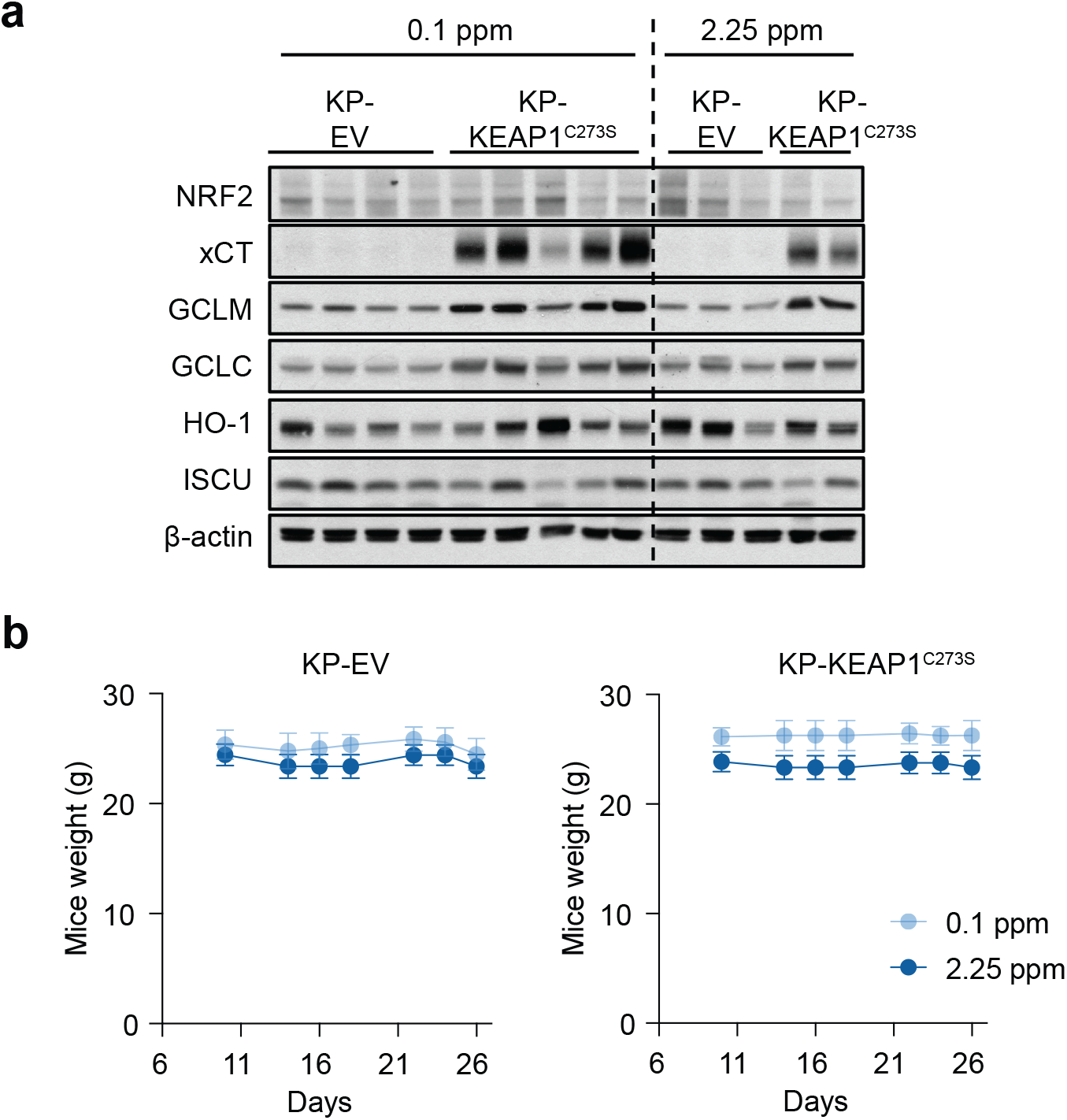
Dietary selenite does not cause weight loss. **a**. Immunoblots of endpoint tumor lysates from the experiment in Fig. 7a**-d**. β-actin, loading control. Each lane is an individual tumor; genotype and diet are indicated above. Lysates were prepared from every tumor yielding sufficient tissue: n = 4 (EV, 0.1 ppm), 5 (KEAP1^C273S^, 0.1 ppm), 3 (EV, 2.25 ppm) and 2 (KEAP1^C273S^, 2.25 ppm) in each group. **b**. Body weight of mice bearing KP-EV (left) or KP-KEAP1^C273S^ (right) allografts, maintained on diets containing 0.1 or 2.25 ppm selenium as sodium selenite, measured from day 6 to day 26. Data are mean ± s.d.

